# A multimodal characterization of the human uncinate fasciculus

**DOI:** 10.64898/2026.03.05.708309

**Authors:** Kelly Perlman, Sarah Barnett-Burns, John Kim, Valérie Pineau Noël, Armand Collin, Justine Major, Malosree Maitra, Anjali Chawla, Murielle Mardenli, Sébastien Jerczynski, Maria Antonietta Davoli, Gabriella Frosi, Julien Cohen-Adad, Daniel Côté, Richard Bazinet, Gustavo Turecki, Corina Nagy, Naguib Mechawar

## Abstract

The uncinate fasciculus (UF) is a hook-shaped long-range association white matter tract that serves to bidirectionally transmit information between the anterior temporal lobe and the orbitofrontal cortex. Neuroimaging studies have suggested that changes in UF microstructure are involved in the neurobiological sequalae of childhood abuse (CA). Given that the UF is not present in rodents, it is vastly understudied with no cellular and molecular information available. To this end, we aimed to perform a multimodal characterization of the UF between individuals diagnosed with depression who died by suicide with (DS-CA) and without a history of severe CA (DS) and psychiatrically healthy individuals (CTRL). Fresh frozen UF tissue was obtained from the Douglas Bell-Canada Brain Bank, with phenotypic information collected via psychological autopsy. Immunohistochemistry with PDGFRα and NogoA was used to label oligodendrocyte precursor cell (OPC) and oligodendrocyte (OL), respectively, and stereology was performed to ascertain cell density and soma volume. Single nucleus RNA sequencing (snRNAseq) was used to generate a transcriptomic survey of the cell types found in the UF. Finally, spectral focusing Coherent Anti-Stokes Raman Scattering (sf-CARS) microscopy was employed in tandem with a custom AxonDeepSeg segmentation model to measure axon diameter, myelin thickness, and g-ratio. No group differences were observed in histology or ultrastructure metrics, but nearly 50 differentially expressed genes (DEG) were identified between groups. Interestingly NECTIN3, the top DEG downregulated in OL1 and OL3 of DS-CA, is a computationally predicted target of the microRNA MIR646, the host gene of which was significantly downregulated in multiple cell types in DS-CA. Age-associated changes were pronounced and observed in all modalities, including an age-related increase in OL density, extensive changes in glial gene expression, as well as decreases in axon diameter and g-ratio.

This study serves as a foundational resource on the molecular and cellular properties of the human UF. Our results suggest that observable myelin-related traces of depression or CA are limited in the UF and highlights the need for future research on the cellular and molecular properties of white matter tracts during aging.

## Introduction

The uncinate fasciculus (UF) is a hook-shaped fiber bundle that transmits information bidirectionally from the anterior temporal lobe to the orbitofrontal cortex, and is considered to be a limbic structure ^1,2^. The temporal terminations of the UF include the temporal pole, the anterior part of the superior, middle, and inferior temporal gyri, and the uncus. Liakos et al (2021) consistently showed the frontal terminations of the main stem of the UF to be the posterior orbital gyrus, pars orbitalis, gyrus rectus, and the posterior part of the paraolfactory gyrus.

With respect to its microstructural properties, the UF has a highly protracted development, and is one of the last fiber tracts to reach full microstructural maturity, with fractional anisotropy peaking after age 30 ^3^. As for function, the UF is not involved in the generation of emotion, motivation or memory itself. Rather, Von Der Heidi et al (2013) posit that its function is thought to lie in a specific intersection of these domains, in which learning from reward and punishment feedback affects the valuation of stimuli and future decision making based on this value. As such, cognitive processes associated with the UF include reversal learning, understanding the higher-level emotional meaning of concepts, and the mediation of approach/avoidance behaviours ^2^.

Most of our knowledge on the UF is derived from neuroimaging studies, particularly diffusion tensor imaging (DTI), in which this tract has been implicated in the neurobiology of a variety of conditions including frontotemporal dementia, epilepsy, psychopathy/acquired criminality, and of particular interest - childhood maltreatment and socioemotional deprivation ^2,4–8^.

Neuroimaging studies have implicated the UF in the neurobiological sequalae of childhood maltreatment, henceforth referred to as childhood abuse (CA). This effect was first identified in a small group of children who underwent severe neglect in Romanian orphanages in the 1980s and 1990s ^4^. Since then, a number of neuroimaging studies on CA and other forms of early-life adversity have demonstrated microstructural changes in the UF ^5,9–14^, though with inconsistent results (i.e., differences reported in significance, laterality, and directionality of findings).

CA is a uniquely strong risk factor for the development of mental illness. In fact, having experienced CA comprises 54% of the population attributable risk (PAR) for depression and 67% of the PAR for suicide attempts ^15^. Furthermore, individuals with a history of CA on average have lower rates of response to treatment in depression and worse overall health outcomes ^15–17^. Importantly, the UF has also been associated with depression in DTI studies ^18^. However, CA and depression are often confounded because few studies of depression reliably measure history of CA. White matter (WM) changes across the cortex have been widely reported in both human and animal studies of early-life adversity, though almost none have evaluated tracts such as the UF. This is notable, as WM changes have been repeatedly associated with early-life stress paradigms in animals ^19–21^ and CA studies in humans ^22^.

From a cellular and molecular perspective, the UF has been greatly understudied because no analogous tract has been identified in rodents, the main animal model in neuroscience ^23–25^. The characterization of human WM tracts is particularly important, as these regions have been increasingly associated with age-related degeneration and termed “hotspots” of aging ^26–28^, Thus, establishing a foundational understanding of the UF in humans is essential to elucidate its involvement in normal and pathological brain functions throughout the human lifespan. This study represents a comprehensive, multimodal characterization of the UF which 1) establishes baseline cellular, molecular, and ultrastructural metrics and 2) assesses the relationships between these measures, CA, depression, and age.

## Methods

### Subject information

In order to disentangle the neurobiological effects of depression from those of CA, we employed a 3 group model: 1) individuals with depression who died by suicide with a history of severe childhood abuse (DS-CA), 2) individuals with depression who died by suicide without a history of childhood abuse (DS), 3) individuals who died naturally or accidentally with no depression nor history of childhood abuse. This research was approved by the Douglas Hospital Research Ethics Board. All brains samples were provided by the Douglas-Bell Canada Brain Bank (www.douglasbrainbank.ca), in partnership with the Québec coroner’s office. The donor’s next of kin provided informed consent upon death, and phenotypic information was obtained via psychological autopsy. This process involved a proxy-based interview with the donor’s next of kin, in which adapted standardized questions were administered, including the SCID-I and II, and the Child Experiences of Care and Abuse (CECA) Interview (Bifulco et al., 1994). CA is defined here as physical, sexual, and/or emotional abuse, or parental neglect before the age of 15 years. Only donors with CECA scores indicating severe CA were added to the DS-CA cohort, and known CA was an exclusion criterion for the DS and CTRL cohorts. Additional information was obtained through the coroner’s report and medical records were supplemented when available. Information such as tissue pH, postmortem interval and refrigeration delay were provided along with toxicological information to measure substances in the body at time of death, and prescription medications for the 3 months prior to death.

### Tissue dissection

The UF was dissected from two locations along its temporal segment by expert brain bank staff in order to obtain enough tissue for all experiments. The dissections were performed using the guidance of a human brain atlas (Mai et al., 2007) on 0.5 cm thick frozen coronal slabs. For the cell density and stereology experiments, blocks of fresh-frozen UF tissue were dissected from the left hemisphere, at the level where the frontal and temporal lobes meet, lateral to the amygdala.

For the remainder of the experiments, fresh-frozen UF was dissected from the left BA38 (temporal pole). The landmarks used included the planum polare, lateral occipitotemporal sulcus, and inferior temporopolar region. These locations were selected because they are close in proximity, and at these locations the UF is not entangled with concurrently running tracts such as the inferior frontal occipital fasciculus. All tissue was stored at-80°C until further processing.

### Cell density and stereology

Subject information for histology experiments can be found in Supplementary Table 1. The UF frozen samples were fixed for 24 hours in 10% formalin at 4°C and then placed in 30% sucrose for a minimum of 24 hours, and until the samples became saturated and sunk. The blocks were sliced into 40 μm thick free-floating sections at-20°C using a Leica CM1950 cryostat.

Immunohistochemistry was performed with antibodies against NogoA (labels mature OLs; Millipore ab5888) and PDGFRα (labels OPCs; R&D Systems AF-307-NA) using 4 sections per subject, with intervals of 6 slices between them (240 μm). After a 10 min incubation in 3% H_2_O_2_ to quench endogenous peroxidase, a 10% blocking buffer containing 0.2% TritonX and normal serum was applied for 1 hour at room temperature and then incubated with primary antibody diluted in blocking buffer (1:500 NogoA; 1:100 PDGFRα, Supplementary Table 2) overnight at room temperature. After a series of 3 washes in PBS, the tissues were incubated in biotinylated goat anti-rabbit secondary antibody for NogoA (Vector Laboratories BA-1000) and biotinylated horse anti-goat secondary antibody for PDGFRα (Vector Laboratories BA-9500) diluted in blocking buffer for 2 hours at room temperature. After another series of washes, the tissue was incubated with Vectastain Elite ABC kit (Vector), which contains streptavidin conjugated to horseradish peroxidase. After a last set of washes, the reagents of the DAB Peroxidase Substrate Kit (Vector) were added to the tissue, such that the HRP would oxidize the DAB into polybenzimidazole, giving the markers a dark brown color such that they could be visualized with light microscopy. Lastly, the sections were mounted onto Superfrost Plus slides and dried overnight at room temperature. Finally, the sections were dehydrated using ascending concentrations of ethanol followed by xylenes and then coverslipped with Permount mounting media. StereoInvestigator v2018.2.1 (MBF Bioscience) was used to perform stereology. OPC/OL density was calculated from measurements taken with the optical fractionator probe (cell quantity) and Cavelieri estimator (tissue volume). Cell body volume was calculated using the nucleator probe and followed by the 2-dimensional cell body area measurement for a final volume.

### UF dissociation and processing

SnRNAseq experiments were performed only on DS-CA and CTRL groups for cost effectiveness purposes, and the workflow is summarized in Figure 2A. The sample size was selected based on a previous postmortem brain study conducted with n=34 ^29^ and which successfully identified nearly 100 depression-associated differentially expressed genes (DEGs). Subject information for the snRNAseq cohort can be found in Supplementary Table 3.

Before beginning the nuclei extraction procedure, all buffers were filtered through 0.22 μm filters to eliminate bacteria, and RNAseInhibitor (Sigma, cat# 3335402001, 1:1000) was added. In each experiment, 4 subjects (2 subjects from each DS-CA and 2 subjects from CTRL) were processed. On average 40-60 mg of tissue was placed in a glass douncer with 3 mL of lysis buffer (5% bovine serum albumin (BSA) wash buffer with 0.05% NP40 detergent). The tissue was dounced 10 times with a thin pestle and 5 times with the thick pestle, and then incubated in the lysis buffer for the remainder of 4 minutes. Next, 10 mL of 5% BSA wash buffer was added to the homogenate, which was all filtered through a 30 μm MACS SmartStrainer (Miltenyi Biotec cat#130-098-463), and spun in a spin bucket centrifuge at 500 g at 4°C for 6 minutes. The supernatant was decanted, the pellet was resuspended in 5mL of 2% BSA wash buffer, filtered through a second 30 μm MACS SmartStrainer, and then spun again at 500 g at 4°C for 6 minutes. The supernatant was decanted, and the pellet was resuspended in 1 mL of 2% BSA wash buffer, then filtered again through a 30 μm MACS SmartStrainer. DRAQ5 (or Hoescht, depending on availability) was added to stain nuclei, and RNaseInhibitor (1:1000) was added to preserve RNA integrity. The nuclei were placed on ice until ready to fluorescence-assisted nuclear sorting (FANS) on a BD FACSAria, at 4°C, using a 130 mm nozzle on “purity” mode.

We gated narrowly on singlets based on forward scatter (particle size) and DRAQ5 fluorescence (see Figure 2B for example), in order to remove as much debris as possible, which serves to effectively minimize ambient RNA ^30^. The singlets were sorted into a tube containing 2% BSA with RNaseInhibitor (1:1000). Wide bore pipette tips were used whenever possible to avoid excessive force on the nuclei.

### Library preparation and sequencing

Nuclei quantity, concentration and purity of was measured on the BD Accuri with DRAQ5 dye, and the nuclei suspension post-sort was visualized under an MBF Zeiss Observer microscope, both in brightfield and fluorescence to assess integrity and purity of nuclei (representative micrograph in Supplementary Figure 1A). After obtaining concentrations, the 4 sorted nuclei samples were pooled, with the volume contributed by each sample calculated to account for 25% of the nuclei in the pool. The pool was filtered one final time through a 30 μm MACS filter, and a final concentration of the pool was taken before nuclei capture with the 10X Chromium system. The 10X Genomics 3’ v3.1 Dual Index Gene Expression chemistry was used for nuclei capture, cDNA synthesis, and library preparation. Concentrations and fragment sizes for cDNA and libraries were measured with High Sensitivity RNA Screen Tape on an Agilent Tape Station. Each library was sequenced on an S4 flow cell on the NovaSeq6000 sequencer at the McGill Genome Center. In order to demultiplex the pooled samples by genotype, DNA was extracted from either brain tissue or blood samples using the QIAamp DNA Mini Kit (Qiagen) and sent for genotyping at Genome Québec. The Illumina Infinium Global Screening Array (GSA v3) + Psych Array was used.

### Bioinformatics

The sequenced libraries were processed with CellRanger version 8.0.1 and analyses were conducted with R version 4.4. A visual depiction of main steps for the bioinformatics workflow can be found in Figure 2C. Even though the nuclei were FANS sorted, which is considered the gold standard for ambient RNA removal, it has been shown that ambient RNA can spur artifactual clustering, especially in immature OLs ^30^. As such, we decided to combine the experimental approach with an *in silico* method. We used CellBender version 0.3 ^31^ to remove ambient RNA and to call cells. Specifically, we used the remove-background command with fpr = 0.000005, learning-rate = 0.00005, posterior-batch-size = 32. This yielded a “clean”.h5 file for downstream use. The alignment files (BAM) derived from CellRanger were then filtered by the barcodes derived provided by CellBender. The data were demultiplexed and de-doubletted by genotype using the Demuxlet and scDblFinder tools in the Demuxafy image, selecting the “AnyDoublets” option upon combining results ^32^. Using “AnyDoublets” indicates that a given nucleus should be flagged as a doublet if it was called as a doublet in either tool. Supplementary Figure 1B contains a representative upset plot showing the Demuxafy results for a single batch.

The data was analyzed using Seurat version 5.0. Seurat objects for each batch/library were filtered to exclude doublets, include genes if they were expressed in at least 3 nuclei, and exclude nuclei who did not express at least 200 genes^33^. Even though we are studying nuclei (not whole cells), there is evidence that nuclei may contain some mitochondrial transcripts due to mitochondrial-nuclear interactions ^34^, but is more likely that mtRNA may be stuck to the nucleus as ambient RNA (due to incomplete removal of all cytoplasmic contents) ^35^ and thus be indicative of a lower quality nuclei. Therefore, nuclei with mitochondrial percent (mtRNA) > 5% were excluded. Data normalization was performed using the sctransform pipeline ^36^, and integration/batch correction was done using Harmony ^37^, with integration performed across batch. The UMAP of nuclei before and after harmony corrected can be seen in Supplementary Figure 1C.

### Cluster annotations

Louvain clustering was performed with a resolution of 0.5 selected as optimal based on visualization from the clustree package ^38^ (Supplementary Figure 1D) and marker gene distribution. Cluster annotation was performed with a combination of manual labelling and label transfer using the Allen Brain Institute’s MapMyCells tool (https://portal.brain-map.org/, RRID:SCR_024672). The manual genes to annotate broad clusters included ALDH1L1 and AQP4 for astrocytes (Astro), MOG and PLP1 for oligodendrocytes (OL), PDGFRA and CSPG4 for oligodendrocyte precursor cells (OPC), CX3CR1 and CSF1R for microglia (Micro), CLDN5 for endothelial cells (Endo), PDGFRB for pericytes, RBFOX3 for neurons, SLC17A7 for excitatory neurons (Excit), and GAD1 for inhibitory neurons (Inhib). Further markers for evaluating excitatory neurons include SATB2, FXDY6, CUX2, GSG1L, and RASGRF2 and for evaluating inhibitory neurons include GAD2, SST, PVALB, VIP, LAMP5, LHX6, ADARB2, CCK, and NPY. To further evaluate the oligodendrocyte-lineage, which includes committed oligodendrocyte precursors (COP), we used MOBP, MAL, OPALIN, CNP, SOX10, OLIG2, BCAS1, PCDH15, VCAN, TCF7L2, TNR, and BCAN. The MapMyCells label transfer was performed with the 10X Human Medial Temporal Gyrus (MTG) SEA-AD dataset using the Deep Generative Mapping algorithm (Supplementary Figure 2A). We then employed tools from the scclusteval package to evaluate cluster concordance across different labelling methods (Supplementary Figure 2C, 2D).

We noticed that the clustering initially did not successfully separate endothelial cells and pericytes, so we manually separated the two cell types based on marker genes and distinct positions along the UMAP using the Seurat CellSelector tool. We annotated 2 clusters as “mixed” indicating that the expression pattern was either mixed across canonical markers or non-specific, thus no cell type label could be attributed to the cluster. The marker genes identified using the FindMarkers function for the Mixed1 cluster were primarily mitochondrial genes, interpreted as a low-quality artifactual cluster and was therefore not included in the downstream analysis. The marker genes identified for Mixed2 were a combination of OPC genes, microglia genes and mitochondrial genes. Importantly, this cluster was small (n=122) and driven by few subjects (Supplementary Figure 3A, 3B). This cluster was also filtered out of the downstream analyses. The distribution of subjects and batch in each cluster is shown in Supplementary Figure 3A and 3B (for individual clusters) and Supplementary Figure 3C and 3D (for broad clusters). The final granular clustering is demonstrated in Figure 1D and broad clustering (aggregation of clusters into broader cell types) is demonstrated in Supplementary Figure 2B.

**Figure 1.**
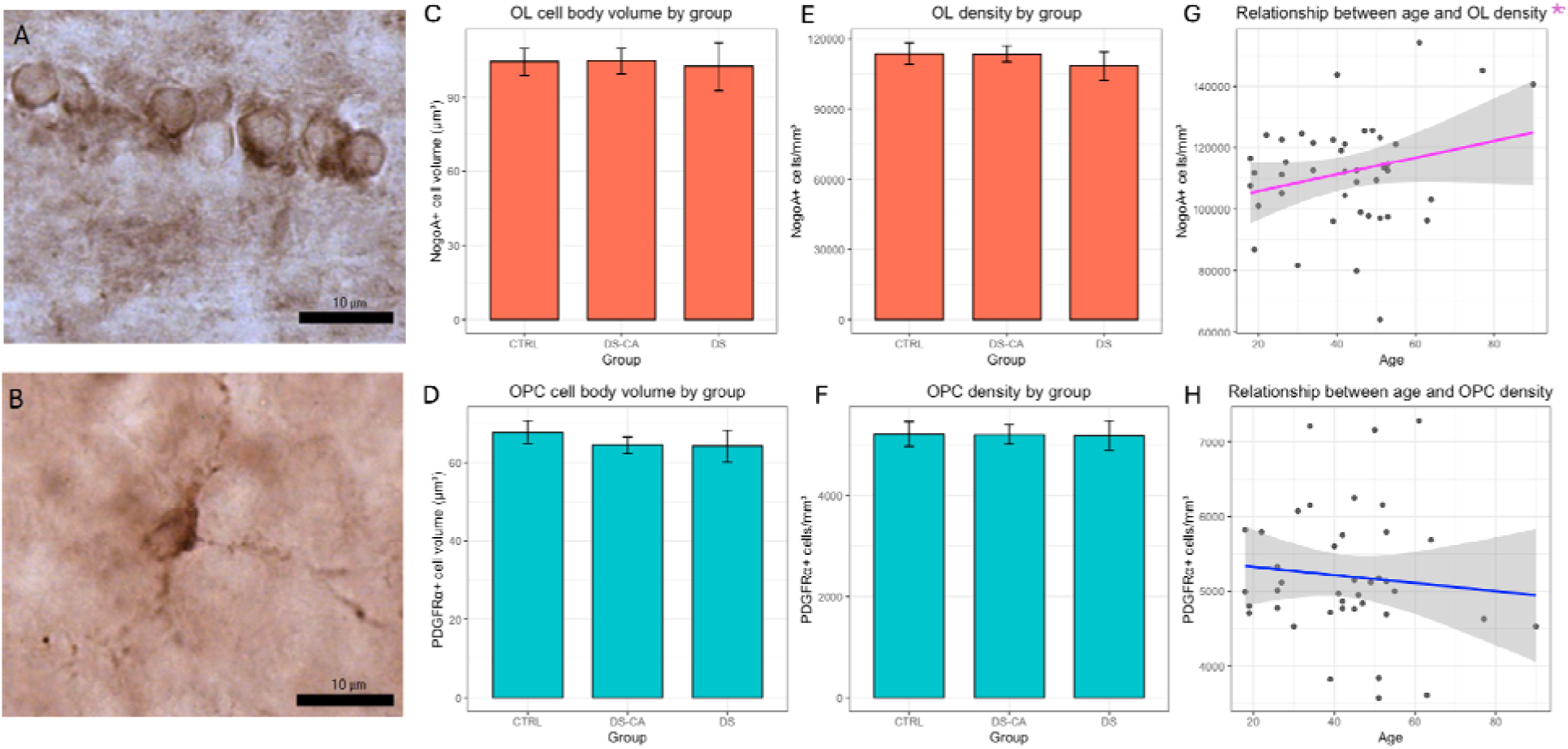
Stereology analysis shows no group differences in OL-lineage cell density or soma volume but shows increasing OL density with age. Representative micrographs of A) NogoA immunoreactive OLs and B) PDGFRα immunoreactive OPCs. Scale bar = 10 µm. Images were published in Dr. John Kim’s thesis ^115^. Bar plots showing groups means ± standard error of the mean for C) OL soma volume and D) OPC soma volume. Bar plots showing groups means ± standard error of the mean for E) OL density and F) OPC density. Scatter plots with trend line showing the relationships between age and G) OL density and H) OPC density. Magenta trend line denotes positive model coefficient for age while blue trend line denotes negative model coefficient for age. Only the age and OL density relationship was statistically significant (p = 0.010).

**Figure 2.**
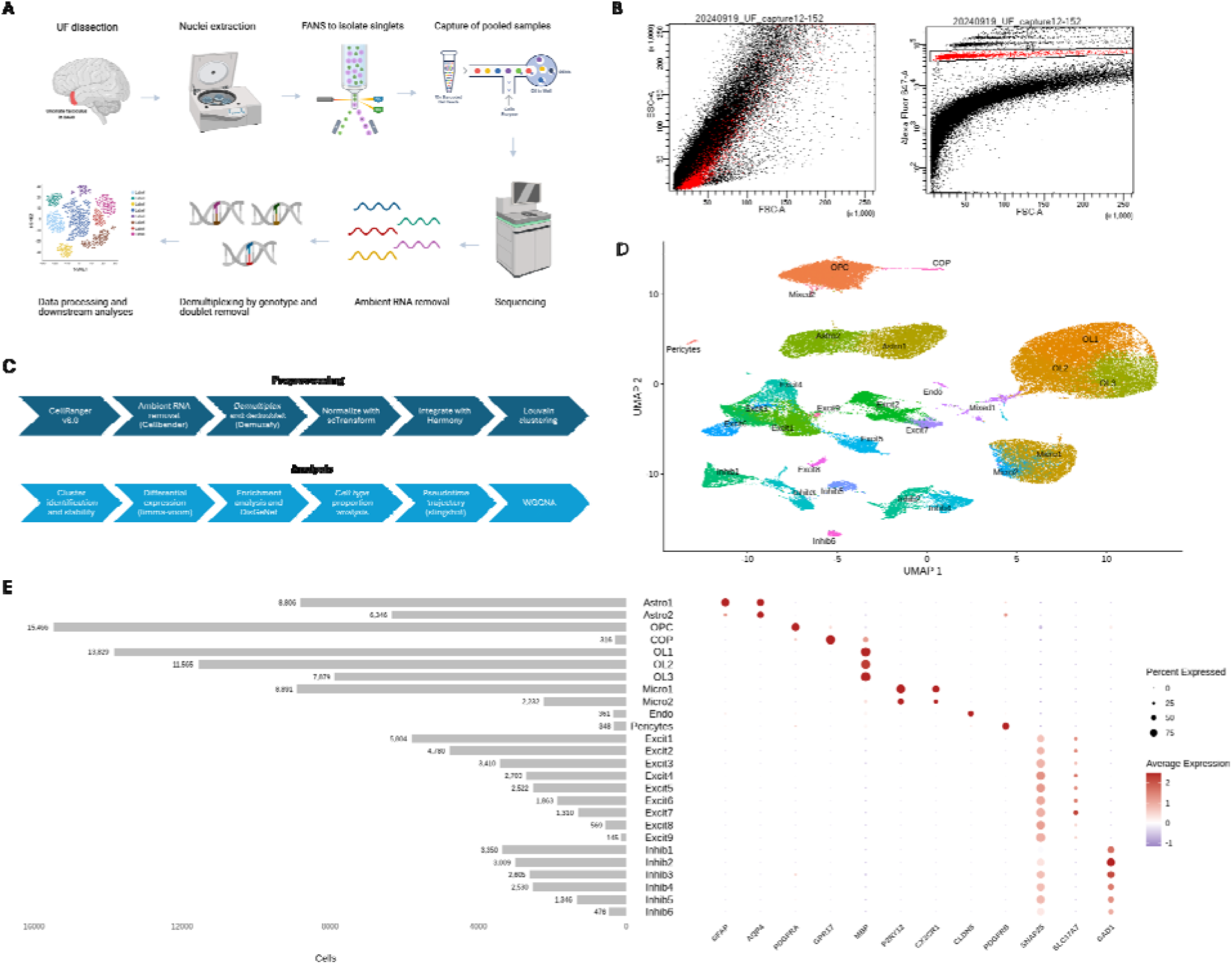
Single nucleus RNA sequencing illustrates cell type landscape of the UF. A) Summary schematic of overall protocol for snRNAseq of the UF. Created with https://BioRender.com. B) Example fluorescence-assisted nuclear sorting (FANS) plot. Singlets were gated in right plot based on forward scatter (FSC) and Alexa 647 fluorescence (from DR dye). The singlets (P1) were then sorted, while debris and multiplets were discarded. Left plot highlights in the sorted singlets in red on a FSC and side scatter (SSC) plot. C) Visual depiction of main steps of snRNAseq bioinformatic workflow. Steps are separated into data preprocessing (top) and analysis (bottom). D) UMAP plot showing nuclei labelled by their individual cell type clusters. E) Plot showing the number of nuclei per cluster and cluster markers genes for cell type characterization. The horizontal bar plot visually depicts the number of nuclei assigned to a given individual cluster and the dot plot shows marker genes for each individual cluster. Mixed clusters were removed from the downstream analyses. Dot plot circles are colored by average expression, and the size of the circle refers to the percent of nuclei in the cluster expressing the marker gene.

**Figure 3.**
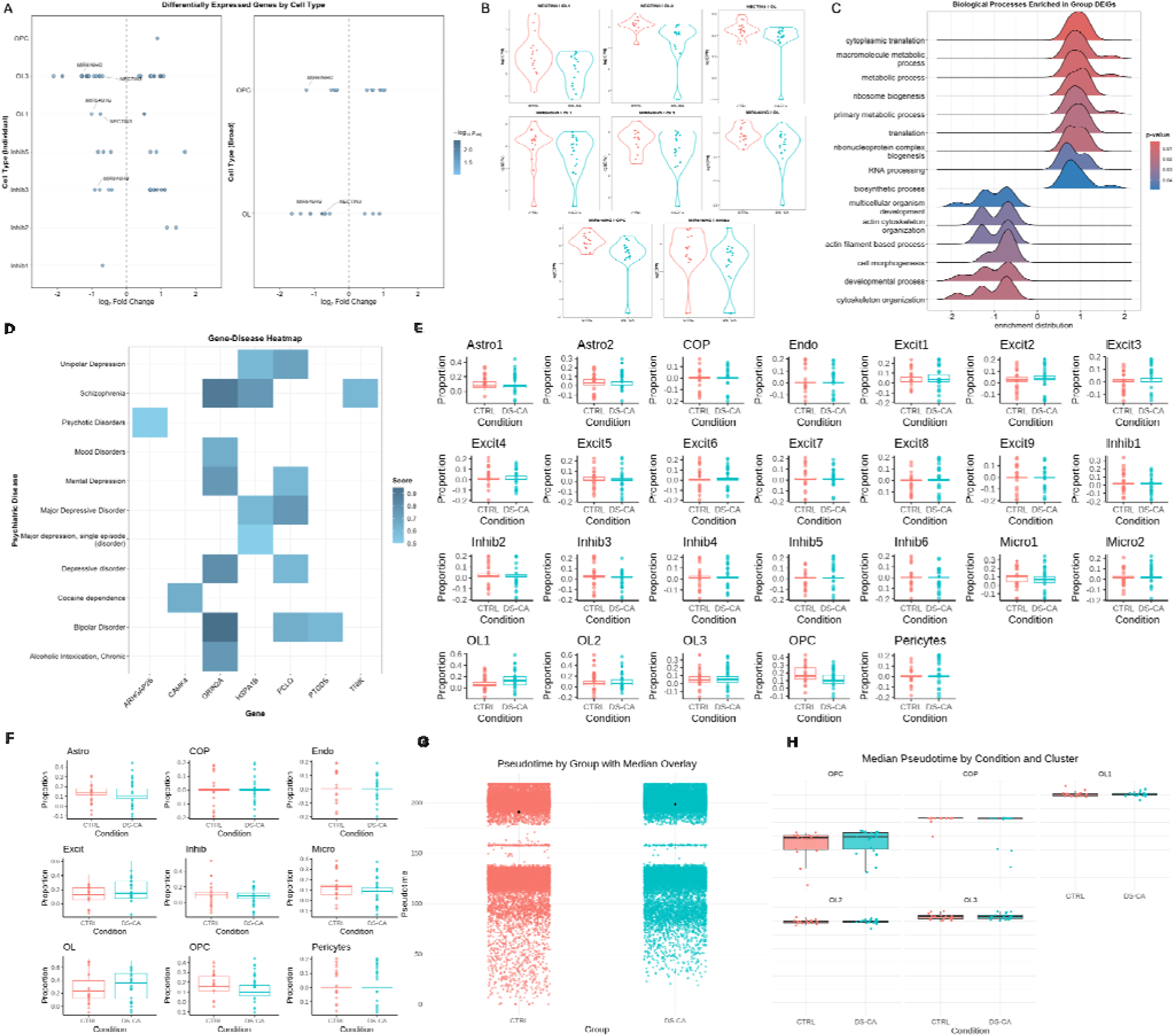
Cell type specific differential gene expression between DS-CA and CTRL. A) DEG summary plot for individual clusters and broad clusters. The x-axis represents the log fold change and y-axis contains the names of clusters containing significant DEGs. The circles are colored by adjusted p-value. The circles corresponding to NECTIN3 and MIR646HG are labeled. B) Violin plots showing normalized pseudobulked expression per subject by group of NECTIN3 and MIR646HG in clusters with p-adjusted < 0.1. Data from subjects present in 2 batches were collapsed via pseudobulk summation for this visualization. Subjects were required to contribute a minimum of 5 nuclei to the individual cluster or 10 nuclei to the broad cluster to be included. C) Ridge plot showing gene set enrichment of group DEGs in the individual clusters considered together. The x-axis represents the enrichment distribution with negative values indicating downregulation in DS-CA and positive values representing upregulation in DS-CA. The y-axis lists the gene sets enriched in the group DEGs. The color of the ridge plot represents the p-value. D) Gene-disease heatmap with the DEG name on the x-axis and psychiatric conditions on the y-axis. The color of the square represents the level of evidence supporting the given gene-disease association, with a minimum score of 0.5 (via DisGeNET). Boxplots showing E) individual cell type and F) broad cell type cluster proportion by group. CTRL data is shown in salmon and DS-CA data is shown in in teal. Mixed clusters are not shown. G) Distribution of OL-lineage cell pseudotime in CTRL subjects (salmon) and DS-CA subjects (teal). The black diamond represents the median pseudotime per group. H) Boxplots showing median pseudotime in CTRL subjects (salmon) and DS-CA subjects (teal) in OPC, COP, OL1, OL2, or OL3 clusters. Clusters are ordered by ascending median pseudotime.

### Cluster stability

In addition to using clustree and label transfer tools, we assessed the quality of clustering by measuring its stability. To do this, we randomly subsampled 75% of the data 100 times and performed clustering using the same resolution (0.5) with each subsample, then calculated the adjusted rand index (ARI) with the existing individual cluster annotations. The mean ARI was 0.848, indicating very high stability across 100 iterations.

### Variance partitioning analysis

To determine which covariates to include in the differential expression model, we performed a variance partitioning analysis with pseudobulked data. Batch explained the most variance of any covariate (Supplementary Figure 4A). Next, we performed a canonical correlation analysis (CCA) to quantify correlation across covariates (Supplementary Figure 4B). Any covariate with a correlation coefficient greater than 0.5 was not included (e.g., PMI, sequencing pool, substance abuse/dependence). It is worth noting that age and batch are moderately correlated (r = 0.44), so in controlling for batch we are underestimating the effects of age, as such any changes with smaller effect sizes might be missed. Group is not correlated with batch (r = 0.08), due to the experimental design of 2 CTRL and 2 DS-CA in each batch.

### Differential expression

Differential expression was performed on pseudobulked objects using the limma package ^39^ with the voom method ^40^, known hereafter as limma-voom. Some subjects are represented in the data twice, because 2 different libraries were generated to increase the number of nuclei obtained per subject. Low-quality nuclei were removed with scater isOutlier(nmads = 3, log = TRUE, type = lower), and genes were retained only if they had >1 count in ≥10 nuclei overall. Gene counts were pseudobulked with the muscat aggregateData function, which summed counts over cluster and subject stratified by batch (i.e., cluster and subject x batch). For each cluster, we included subjects contributing ≥ 5 nuclei (individual clusters) or ≥ 10 nuclei (broad clusters) to ensure sufficient representation. We constructed an edgeR DGEList object, filtered lowly expressed genes using the filterByExpr function, and applied TMM normalization. Models were fit with limma-voom using voomWithQualityWeights = TRUE. To account for subjects present in two batches, we used the following pipeline: 1) apply voomWithQualityWeights without accounting for the within-subject correlations, 2) calculate duplicateCorrelation (blocking on subject ID), 3) apply a final voomWithQualityWeights and lmFit that takes into account the within-subject correlation via the duplicateCorrelation estimates. The group contrast used design ∼ 0 + Group + Age + Sex + pH + Batch. The age effect used ∼ z-scored age + Group + Sex + pH + Batch, in which age was centered and z-scored across all subjects. Statistics were moderated with eBayes (trend = TRUE, robust = TRUE). P-values were corrected for multiple comparisons with the Benjamini-Hochberg method. Of note, the unknown pH for R22_517 was imputed with the average of all pH values for the cohort.

DEG with adjusted p-value < 0.1 were considered as significant, as in ^29^. For group DEGs, log fold change (logFC) represents the log base 2 difference between DS-CA and CTRL, in which positive logFC indicates an upregulation of the gene in DS-CA compared to CTRL, and negative logFC indicates a downregulation of the gene in DS-CA compared to CTRL. For age DEG, the logFC value represents the slope of expression change for 1 standard deviation increase in age (in log base 2 scale). Positive logFC represents increase with age and negative logFC represents decrease with age.

To ensure that the DEGs were not artifactual driven by low numbers of subjects, we added the following filters: for group DEGs, the gene was required to be expressed (i.e., not NA and > 0) in more than 75% of subjects from each group (DS-CA n ≥ 12, CTRL n ≥ 15), and for age, DEGs were required to be expressed in more 75% of all subjects (n = 27). These filters were applied for both broad and individual clusters. The gseGO function from clusterProfiler was used to characterize gene set enrichment within the differentially expressed genes, with the following parameters: ont= “ALL”, keyType = “ENSEMBL”, nPerm = 10000, minGSSize = 10, maxGSSize = 500, pvalueCutoff = 0.05, verbose = TRUE, OrgDb = organism, pAdjustMethod = “none”.

### Cell type proportions

Proportions of nuclei in each cell type (both broad and individual clusters) was compared between groups using the protocol described in ^41^. For each subject, the number of nuclei in each cluster was divided by the total number of nuclei (for both bulk and individual clusters) to calculate cluster proportions. Then, proportions were compared between DS-CA and CTRL using a two-sided Wilcoxin test. To reduce the influence of potential outliers, each test was bootstrapped 10000 times.

### Gene-disease associations

Gene-disease associations (GDA) were investigated with DisGeNet, using the disgenet2r R package (https://gitlab.com/medbio/disgenet2r), querying the PsyGeNet database with GDA scores between 0.5-1 as accepted. GDA scores represent level of evidence for a given association, with 0 representing no evidence for an association and 1 representing maximum evidence for an association.

### Pseudotime trajectory

Pseudotime trajectory analysis on the OL-lineage cells only (includes OPC, COP, and OL broad clusters) was performed with slingshot ^42^. The raw merged Seurat object was subset by the OL-lineage cell clusters and was then normalized with sctransform and integrated by batch with harmony. The UMAP was re-calculated with 15 PC dimensions, and slingshot was run with the PCA as reducedDim, broad OL clusters as clusterLabels, with OPC as start.clus, stretch = 0.1, shrink = 0.7, and reweight = TRUE. The nuclei were annotated with the same OL-lineage individual level cluster labels as the full object (OPC, COP, OL1, OL2, OL3). Pseudotime values overall and in each cluster were then compared by group with the following formula: lmer(pseudotime ∼ Group + Age + Sex + pH + (1|SubjectID)).

### Overlap with mouse OL aging dataset

To establish whether there exists an overlap in the aging fingerprint of OLs across species (mouse and human), we leveraged the aging dataset by Ximerakis et al, specifically the OLG cluster from Supplementary Table 6 ^43^. Using the MGI Mouse–Human Homology resource (from https://www.informatics.jax.org/homology.shtml), we removed all mouse-specific genes (genes for which there is no human ortholog) from the Ximerakis dataset and all human specific genes (genes for which there is no mouse ortholog) from the present OL cluster datasets (broad and individual clusters) to ensure a comparable gene list across datasets. We then took all the genes significantly associated with age (adjusted p-value < 0.1 in each dataset) and assessed overlap between the datasets. To quantity whether there was a statistically significant enrichment of overlapping genes, we applied the one-sided Fisher’s exact test (hypergeometric test), separately for the broad OL cluster overlap and the individual OL clusters overlap.

### Pseudobulk Weighted Gene Co-expression Network Analysis (WGCNA)

To better understand the underlying structure of the snRNAseq data and evaluate gene co-expression networks, we employed WGCNA with pseudobulked data as performed in ^41^. WCGNA does not support repeated measures or random effects, so we aggregated counts derived from the cell type of interest (OL) by subject (without batch) using AggregateExpression in Seurat. As such, the counts for subjects represented in 2 batches were aggregated together via summation. Covariates (pH and PMI) were corrected for using limma’s RemoveBatchEffect function, in which covariates to remove can be specified while preserving variables of interest (in this case, age, sex, and group). Genes with low overall counts (less than 5) were removed.

The pseudobulked counts were converted to a DESeq2 object and normalized and transformed with the rlog function. After testing soft powers, a value of 16 was determined to be optimal (Supplementary Figure 8B, 8C) and a minimum module size of 30 genes was a requirement for network construction. After the networks were created, we proceeded to use module trait correlation analysis. Genes that comprised modules significantly associated with group or age were assessed for biological process enrichment with gene set enrichment analysis using the fgsea package. A minimum gene set size of 10 and a maximum gene set size of 500 were used as thresholds for set inclusion.

### Spectral Focusing Coherent Anti-Stokes Raman Scattering microscopy

Tissue ultrastructure was imaged using spectral focusing Coherent Anti-Stokes Raman Scattering (sf-CARS) microscopy, a label-free technique that exploits the lipid-rich nature of myelin by probing the CH_2_ molecule stretching vibrational band (2,845 cm^-1^), which are enriched in fatty acid tails ^44–46^. The subject information for this experiment can be found in Supplementary Table 4. Briefly, fresh frozen UF blocks were fixed overnight at 4°C in 10% formalin, and subsequently kept in 1x PBS at 4°C. The tissue was then sliced in ice cold 1x PBS into 300 mm thick sections on a Leica VT1200S vibratome, and stored in cryopreservative at-20 °C.

The sf-CARS imaging was conducted at Université Laval (CERVO), with the setup including a ytterbium (Yb) Insight X3 pulsed femtosecond laser system with two output beams and a 19.6 cm fused silica rod for temporal chirping (full configuration described in sf-CARS methods paper ^47^). Image acquisition was performed with an Olympus 60X/0.8 NA water immersion objective with an additional 2X digital zoom. A full description of the sf-CARS methodology can be found in ^47^ including the optical imaging parameters and details of the AxonDeepSeg deep learning image segmentation model.

To segment structures (axon and myelin), we trained a custom deep learning model with 2D U-Net architecture using the AxonDeepSeg ^48^ pipeline. The trained model is available for public use: https://github.com/axondeepseg/model-seg-human-brain-cars. The iterative training process involved multiple rounds of manual correction of model predictions, and the final training dataset contained 6977 axons across 124 images. The quality control metrics are detailed in the methods paper ^47^. Imaging resolution was 0.167 um/pixel, so we excluded any axon with a diameter less than 0.335 um, because that would represent a structure less than 2 pixels, which is prone to inaccurate measurements. We also removed axons with eccentricity values > 0.9, as this may represent a highly ellipsoid shape, and the quantification metrics are computed with the assumption that axons are roughly circular in shape.

## Results

### No cell density or volume changes between groups but higher OL density with age

We used immunohistochemistry to stain OL and OPC with well-known markers, NogoA and PDGFRα, respectively (n=42 for both cell types). Representative micrographs are shown in Figure 1A (OLs) and Figure 1B (OPC). Of note, OL processes are not visible with this staining due to the localization of NogoA protein predominantly to the endoplasmic reticulum ^49^. Then, a stereological approach was used to calculate OL and OPC soma volume and density. The mean soma volume of OL was 103.95 ± 3.91 μm^3^ and of OPC was 65.32 ± 1.68 μm^3^. The mean density of OL was 111995.70 ± 2702.65 cells/mm^3^ and of OPC was 5203.44 ± 135.07 cells/mm^3^. For OL, no significant differences between groups were found in soma volume (p > 0.99, Figure 1C) or cell density (p = 0.15, Figure 1E). Likewise, for OPC, no significant differences between DS-CA, DS, and CTRL groups in soma volume (p =0.66, Figure 1D) or cell density (p =0.91, Figure 1F) were observed. No significant relationships were observed between OL or OPC soma volume and age (OL p = 0.55, OPC p = 0.86, Supplementary Figure 5). However, OL density was significantly associated with age (p = 0.010, Figure 1G) in a positive linear relationship. No significant associations with age and OPC density were found (P =0.39, Figure 1H). In summary, we did not see any significant relationship between OL-lineage cell soma volume nor cell density to CA or depression. However, we did observe that OL density was significantly increased as a function of age.

### Cell type specific characterization of the UF with snRNAseq

Figure 2A illustrates the main steps of the snRNAseq workflow. After demultiplexing and doublet removal, we retained 113,503 nuclei from 35 subjects. Unsupervised graph-based clustering identified 28 distinct clusters (Figure 2D), derived from 10 broad cell types (Supplementary Figure 2B), though the mixed clusters were not considered for downstream analysis (see Methods). These nuclei were deeply sequenced, with an average of 89.7% sequencing saturation across libraries, in which saturation is calculated as *1 - (number of valid and deduplicated reads / number of valid reads)*. Sequencing statistics from cellranger and cellbender tools are listed in Supplementary Table 5. Basic marker genes for each cell type and the number of nuclei per cluster are plotted in Figure 2E. A complete list of cell type markers for individual clusters can be found in Supplementary File 1. The number of nuclei per individual and broad cluster are summarized in table format in Supplementary Tables 6 and 7, respectively. The MapMyCells tool was used to map the 10x Human MTG SEA-AD taxonomy onto our clusters (Supplementary Figure 2A). We found overall strong concordance across datasets (Supplementary Figure 2C, 2D), with the adjusted rand similarity index (ARI) between the MTG SEA-AD clusters and our individual clusters at 0.615 and the ARI between the MTG SEA-AD clusters and our broad clusters at 0.821.

Our proportions of broad cell types appeared roughly similar to another snRNAseq WM analysis, with a similar dissociation method ^50^. The percentage of total nuclei that were labelled as neurons in this previous dataset was 28.3%, which is similar to that of the current dataset, at 32.1%.

While we expected some grey matter contamination due to the technical challenge of frozen dissection, we can be confident that at least a substantial portion of these are indeed proper WM neurons (also known as interstitial neurons), as we have recently shown that both the RNA-and protein-level, neuronal cell bodies exist in the UF proper ^51^. We showed that both excitatory and inhibitory neurons exist in the UF at the same dissection location (BA38), as well as a second UF location along the temporal segment ^51^. In the current study, we observed that every neuronal cluster showed a high similarity to a single neuronal cluster from Allen MTG SEA-AD (Supplementary Figure 2C, Supplementary Table 8).

The excitatory clusters in Allen MTG SEA-AD are annotated to a specific cortical layer. Therefore, while all our excitatory neuron clusters showed transcriptomic similarity with cortical neurons, they are derived (at least in part) from the actual WM. Interestingly, 7 of 9 excitatory clusters show most similarity with deep layer neurons (layers 5 and 6), which are more proximal to the WM. It is thus possible that WMN may be more transcriptomically similar to their more proximal cortical neurons as opposed to ones that are more distal. Importantly, we posit that neurons were most robust to our specific tissue dissociation methods used as compared to glia, a phenomenon which has previously been reported and appears to be dissociation method-dependent ^52^.

In order to determine the relative maturity level of nuclei within the OL-lineage, we performed a pseudotime trajectory analysis with slingshot. As demonstrated in Supplementary Figure 6A, the OPC cluster has the lowest median pseudotime, followed by COP, followed by the OL clusters. Among the OLs, OL2 is the least mature, followed by OL1, with OL3 being the most mature.

### NECTIN3 downregulation in OL of DS-CA

Differential expression for group was performed with limma-voom, with covariates age, sex, pH, and batch, conducted for both individual and broad clusters. Samples were pseudobulked by cluster and subject by batch (Supplementary Figure 7). With adjusted p-value < 0.1, we identified 48 DEGs in individual cell types and 22 DEGs in broad cell types (Figure 3A). Of the 48 individual cluster DEGs, 21 were derived from OL3, the most mature OL cluster according to pseudotime. Notably, the calcium independent immunoglobulin-like cell adhesion molecule NECTIN3 was downregulated in DS-CA vs CTRL in OL1, OL3, and OL (broad) (Figure 3B).

Intriguingly, MIR646HG, the host gene for the microRNA MIR646, was significantly downregulated in OL1, OL3, Inhib3, OPC (broad), and OL (broad) (Figure 3B). Gene set enrichment analysis of the individual cluster DEGs showed general terms, including translation, metabolic process, and cytoskeletal organization (Figure 3C). Differential expression results can be found in Supplementary Table 9 (individual cell types, full results in Supplementary File 2) and Supplementary Table 10 (broad cell types, full results in Supplementary File 3). Given the prominence of MIR646HG in the results, we asked whether there was any overlap between the predicted targets of MIR646 and the other significant group DEGs. We sourced a list of high-confidence computationally predicted MIR646 gene targets from the molecular signatures GSEA database (https://www.gsea-msigdb.org/gsea/msigdb/human/geneset/MIR646.html).

Remarkably, observed that two of our DEGs, NECTIN3 and TRIM36, were indeed targets of MIR646, pointing to a potential regulatory mechanism. According to the miRDB database (https://mirdb.org/index.html), both of these genes have a high target confidence score for MIR646 (NECTIN3 score = 92/100 and TRIM36 score = 89/100), indicating that both are strong computationally predicted targets, though still require experimental validation.

To examine whether the protein-coding DEGs have previously been associated with psychiatric disorders, we queried the PsyGeNet database via the DisGeNeT tool with a minimum evidence score of 0.5 required. We identified 4 genes in the individual cluster DEGs (PCLO, ARHGAP26, HSPA1B, TNIK) and 3 genes in the broad cluster DEGs (GRIN2A, PTGDS, CAMK4) that have been linked to psychiatric disease and are summarized in a heatmap colored by GDA association evidence level (Figure 3D). Furthermore, we do not observe any significant differences in cell type proportions between groups, in either individual (Figure 3E) or broad (Figure 3F) cell type clusters.

We then compared the pseudotime of OL-lineage clusters between groups (Supplementary Figure 6b contains a UMAP colored by pseudotime). Though descriptively DS-CA OL-lineage cells seem to be shifted towards a higher pseudotime as demonstrated by Figure 3G, this difference was not significant (p = 0.15). Likewise, no significant group differences in pseudotime were observed at the cluster level in OPC, COP, OL1, OL2, or OL3 (Figure 3H).

Therefore, we do not detect clear shifts in maturity within the OL-lineage between groups.

In summary, a relatively limited number of DEGs were observed between DS-CA and CTRL, but two DEGs, NECTIN3 and TRIM36, were predicted to be targets of MIR646, derived from MIR646HG, itself a DEG. No robust group differences were identified in the cluster proportion or pseudotime trajectory analyses.

*UF glia show gene expression changes with aging and correlate with decreased myelination* In all analyses, we considered age as a continuous variable, including both DS-CA and CTRL, spanning ages 25 to 66 in the snRNAseq dataset. Of the aging DEGs in the individual clusters, 122 DEGs were detected, with all of them derived from glial cells, primarily OL1 (Figure 4A), the second most mature OL cluster according to pseudotime (Supplementary Figure 6A). Of the broad age DEGs, 117 genes were detected, all from glial cells, with the majority from OL (Figure 4B). Differential expression results can be found in Supplementary Table 11 (individual clusters, full results in Supplementary File 4), and Supplementary Table 12 (broad clusters, full results in Supplementary File 5). Unexpectedly, in the individual clusters we noticed that enriched processes were often linked with neuronal development, neurogenesis, and other related terms (Figure 4C). Upon examining the genes in these sets, we noticed that they frequently represented what are canonically considered to be axon guidance or neurogenesis-related molecules (e.g., DCC, PLXNB1, SDK1, RELN, FEZ1), but all derived from OL. In the broad clusters, processes enriched in the DEGs include transporter activity, ion binding, and transmembrane transport (Supplementary Figure 8A).

**Figure 4.**
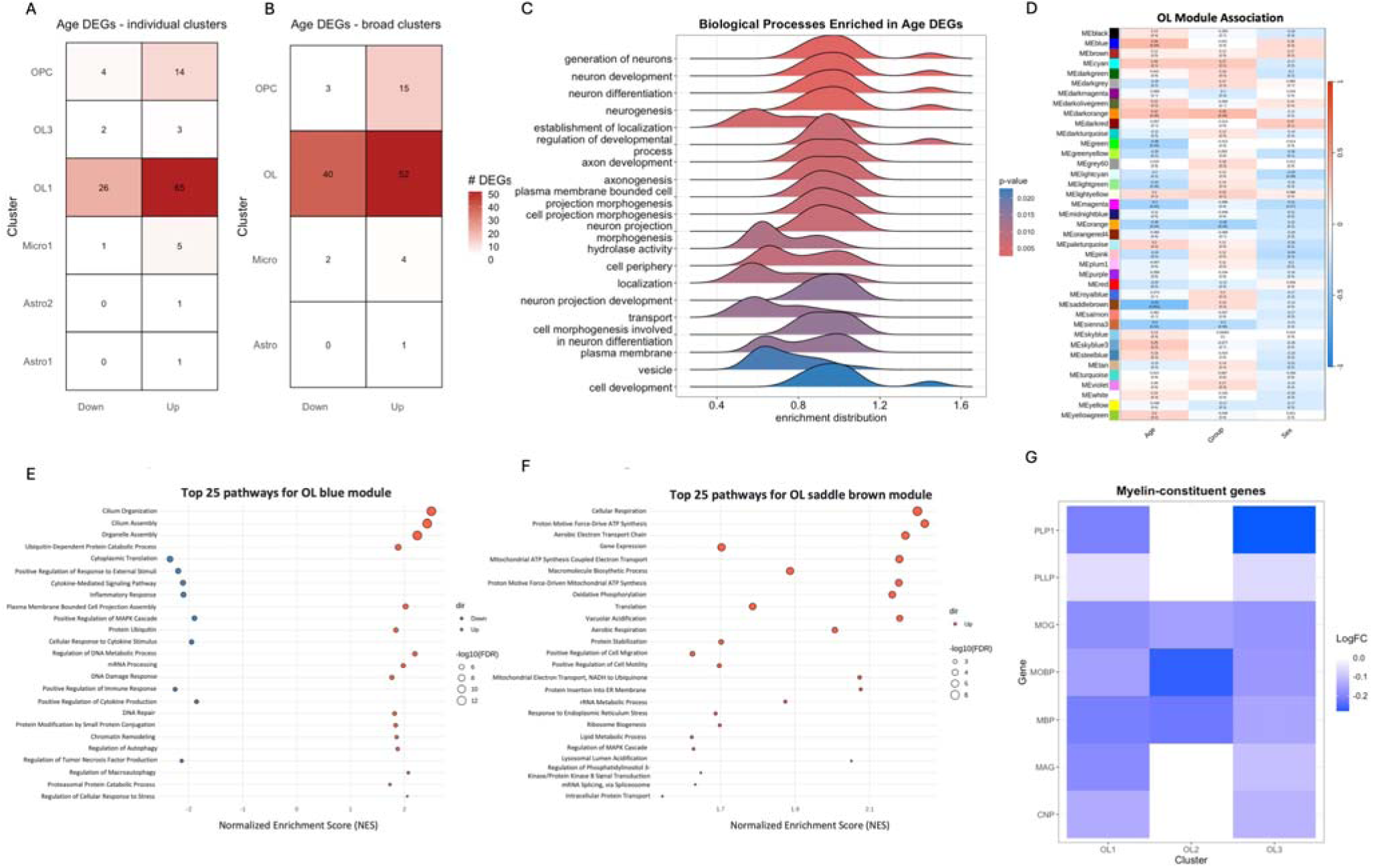
Glial cells show pronounced gene expression changes across the age span. Heatmap displaying number of genes in each A) individual and B) broad clusters that showed statistical significance (adjusted p-value < 0.1) and stratified by down-or up-regulation according to negative and positive log fold change values, respectively. C) Ridge plot showing top 20 gene sets enriched in the aging DEGs in the individual clusters considered together. The x-axis represents the enrichment distribution with negative values indicating a decrease with age and positive values representing an increase with age. The y-axis lists the gene sets enriched in the aging DEGs. The color of the ridge plot represents the p-value. D) Heatmap of WGCNA modules constructed for the OL broad cluster with soft power set at 16 and minimum module size set at 30 genes. Each module is represented by a color shown on the left-hand side of the plot, along with the correlation of each module with age, group, and sex. The correlation coefficient is shown on top of the unadjusted p-values in parentheses. The color bar on the right maps onto the correlation coefficient from-1 to 1, with negative values in blue and positive values in red. The effects of PMI and pH were removed from the normalized gene expression data prior to module construction. Dot plot showing top 25 enriched biological pathways for the genes in the E) OL blue module and F) OL saddle brown module. On the dot plots, normalized enrichment scores (NES) are represented on the x-axis and gene set names on the y-axis. The size of the circle represents the-log10(false discovery rate) value, with larger dots having lower FDR values. The color of the dot represents the direction of regulation of each term (blue = down, salmon = up). G) Heatmap of the logFC of myelin constituent-genes in each individual OL cluster from the age DEG analysis. The color bar represents the logFC value with more negative values (indicating decreased expression with age) colored in darker blue. Expression values are decreased with age in all genes in OL1 and OL3. OL2 shows either decreases or no change. The star on MBP in OL1 indicates p < 0.1 in the age DEG analysis.

Given the high proportion of aging DEGs deriving from OL, we asked whether the current UF aging OL signature would overlap with previously published rodent OL aging signatures. We compared the DEG with adjusted p-value <0.1 in the current dataset with genes differentially expressed in the OL cluster between young and old mice with adjusted p-value <0.1 ^43^. As listed in Supplementary Table 13, we identified 15 genes overlapping as aging DEGs in individual OL clusters (Fisher’s exact test p = 0.46) and 24 genes were overlapping in the broad OL cluster (Fisher’s exact test p = 0.0094), for a total of 30 unique overlapping genes. These results indicate a substantial overlap in the OL aging signature between species, and significantly so when considering human OLs as a broad cluster.

We aimed to further characterize the patterns of gene expression in OL by performing WGCNA analysis to assess whether certain gene co-expression patterns could be linked to age, group, or sex (Figure 4D). We then elected to perform gene set enrichment analysis on the modules with p < 0.05 that had the highest magnitude negative and positive correlations with age. The blue module showed the highest magnitude positive correlation with age (r = 0.34, p = 0.04, Figure 4D), and enriched terms included inflammatory response, cytokine-mediated signalling pathway, protein ubiquitination, DNA repair, regulation of autophagy, and regulation of cellular response to stress (Figure 4E). The saddle brown module showed the highest magnitude of negative correlation with age (r =-0.53, p = 0.001, Figure 4D), and enriched terms included cellular respiration, electron transport chain/oxidative phosphorylation, ribosome biogenesis, chromatin remodelling, response to ER stress, and interestingly, lipid metabolic process (Figure 4F).

Importantly, when considering the top 40 enriched gene sets in the saddle brown module, regulation of myelination is listed consistent with our previous observations (Supplementary Figure 8D). In summary, the processes affected with age include metabolism (oxidative phosphorylation, lipid metabolism), inflammation, transcription/translation (alternative splicing, ribosomal biogenesis) response to cellular stress (ER stress response DNA damage/repair), protein degradation (ubiquination, autophagy), and myelination.

There were two modules for which group was significant (orange and dark orange modules). In the orange module, the expression of which was significantly downregulated for both group (r =-0.39, p = 0.02) and age (r =-0.35, p = 0.04), enriched terms included translation, calcium ion transmembrane import/calcium ion import, and proteoglycan metabolic process (Supplementary Figure 8E). Other terms overlapped with the previously listed module terms (cellular respiration, cytokine production, ribosome biogenesis/translation). The dark orange module, the expression of which was significantly upregulated in group (p = 0.04, r=0.35), and approaching significance in age (p = 0.06, r = 0.32) showed a small number of enriched terms, including chromatin remodelling, axon development, and cell migration (Supplementary Figure 8F). Sex did not show any significant modules in the WGCNA analysis.

As we recently reported, aging in the UF is associated with decreased levels of myelin-constituent genes and proteins ^53^. As such, we plotted the logFC of these genes (MBP, MOBP, MOG, MAG, CNP, PLP1, PLLP) explicitly to assess possible relationships with age (Figure 4G). In OL1 and OL3, all myelin-constituent genes showed negative logFCs, with MBP in OL1 having p-adj = 0.043. In OL2, the genes either had negative logFC or no change. NKX6-2, while not a myelin-constituent gene, is a key transcription factor in the regulation of myelination and was also downregulated with age (p-adjusted = 0.016). In the broad OL cluster, MBP was significantly downregulated as well (p-adjusted < 0.1). Taken together with the results from our previous work, the evidence indicates that in the UF, the expression of myelin-constituent genes declines with age ^53^.

In summary, a wide variety of biological processes, both myelin-specific and more global, are dysregulated with age in the UF. All the aging DEGs were derived from glia, with the most derived from OLs, and the OL aging signature overlapped significantly with mouse OLs in a previously published aging dataset ^43^.

### Myelin ultrastructure changes associated with age, not group

To measure UF myelin ultrastructure, we implemented sf-CARS microscopy on UF samples (n = 30, Supplementary Table 4). Axon and myelin structures were segmented using a custom trained AxonDeepSeg model, now available for public use (https://github.com/axondeepseg/model-seg-human-brain-cars). An example segmentation can be seen in Figure 5A. After quality control filtering as described in Methods, a total of 18,822 axons were detected, with their binned size distribution demonstrated in Figure 5B. Across all axons, the mean axon diameter was 0.95 ± 0.56 µm, the mean myelin thickness was 0.49 ± 0.15 µm, the mean g-ratio was 0.48 ± 0.095. The mean and standard deviation of each of these metrics stratified by diameter bin is represented in Supplementary Figure 9A.

**Figure 5.**
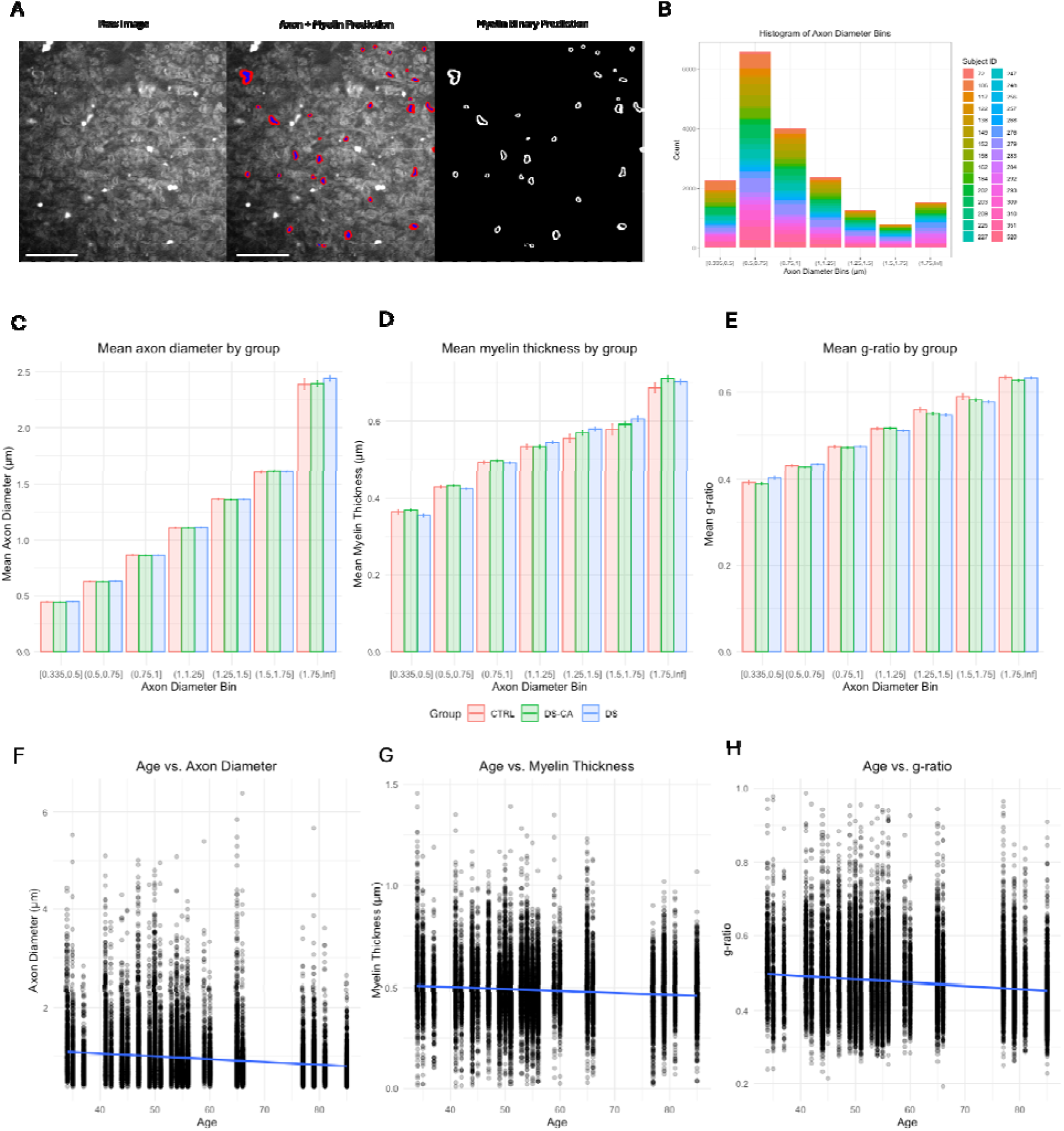
Spectral Focusing Coherent Anti-Stokes Raman Scattering reveals ultrastructural changes with aging. A) Micrograph of representative sf-CARS image (left), the segmentation of axons and myelin from that image (middle), and a binary segmentation of the myelin (right). Scale bars = 20 µm. B) Histogram showing frequency of axon per diameter bin. The histogram is colored by the subject ID, showing that each subject shows a distribution of axon diameters, and there are no diameter bins driven by a particular subject or set of subjects. Bar plot showing mean ± standard error of the mean per group for C) mean axon diameter, D) mean myelin thickness, and E) mean g-ratio across axon diameter bins. CTRL data is shown in pink, DS-CA data is shown in green, and DS data is shown in blue. Scatter plots with trend line (blue) showing the relationship between age and F) axon diameter, G) myelin thickness, and H) g-ratio. All metrics decrease with age, but axon diameter and g-ratio decrease significantly (p = 0.039, p = 0.017, respectively).

To assess the relationship between ultrastructure and lipid composition, we leveraged our extant UF lipid concentration data ^53^. Interestingly, the concentrations of the most abundant phospholipids in myelin (ethanolamine glycerophospholipids and choline glycerophospholipids) as well as the overall total UF lipid concentration were significantly positively correlated with ultrastructure metrics. The cholesterol concentration showed a positive but non-significant correlation with ultrastructure metrics (Supplementary Figure 9B).

When considering all the axons together, no significant effect of group was found for any metric (Supplementary Figure 9C), which held true when stratifying by axon diameter (Figure 5C).

However, as demonstrated in Figure 5D, significant inverse relationships were observed between age and axon diameter (p = 0.039) and between age and g-ratio (p = 0.017), while a non-statistically significant inverse relationship was observed for age and myelin thickness (p = 0.24). When stratifying by axon diameter bin, statistical significance for age was not detected (despite a trend with p = 0.0599 for axon diameter in the 1-1.25 µm bin) likely due to the decrease in power that results from the stratification (Supplementary Figure 9D).

Furthermore, we observed no significant effect of group or age on fiber density (Supplementary Figure 10A, 10B). However, as demonstrated in Supplementary Figure 10C and 10D, the density patterns of small (less than 1.5 µm) and large fibers (great than 1.5 µm) were divergent. With age, the density of small fibers was increased, while the density of large fibers was decreased. It is important to note that the fiber density only represents myelinated axons, as sf-CARS is not tuned to capture non-myelinated axons.

In summary, none of the measured UF ultrastructural parameters show a relationship with CA or depression. However, axon diameter and g-ratio showed significant decreases with age.

Therefore, it appears that thinner axons predominate in aging, possibly at the expense of thicker axons, at least in this WM tract.

## Discussion

The multimodal characterization of human postmortem UF described in this paper represents an important advancement in our understanding of human WM. We provide the first ever snRNAseq dataset for the UF, characterizing the array of cell types and providing transcriptomic evidence for the presence of WM neurons recently described in this tract ^51^. We additionally provide histological metrics of OPC and OL, and an ultrastructural characterization of myelinated fibers using a novel sf-CARS and deep learning segmentation approach. We then employed these methods to inquire about the effects of CA, depression, and aging in a multimodal fashion.

The top DEG in OL was NECTIN3 (also a top DEG in OL1 and OL3), which was downregulated in DS-CA compared to CTRL. NECTIN3 is a calcium-independent, immunoglobin-like cell adhesion molecule expressed primarily at the post-synaptic neuron ^54^. There is a substantial body of literature showing Nectin3 downregulation in the mouse brain ^54–58^ in response to early life and adulthood stress, thus impairing synaptic adhesion. In fact, early life adversity causally downregulates hippocampal Nectin3 levels in a corticotrophin releasing factor-dependent fashion, with the decrease in Nectin3 mediating cognitive impairments and dendritic spine loss. In a chronic restraint stress model, Nectin3 was downregulated in the CA1 region of the mouse hippocampus. Matrix metalloproteinase 9 (Mmp9) showed increased gelatinase activity in response chronic stress, and it is known that Mmp9 cleaves Nectin3 ^58^. This result is of great interest given that changes in the extracellular matrix, specifically in the form of perineuronal nets (PNN), have been identified as a possible mechanism underlying the neurobiology of early life adversity and stress ^59–65^. However, NECTIN3 has not been investigated from an OL-lineage or myelination standpoint, and how that may be linked to stress. Given the very low density of neurons in the UF ^51^, we would not expect many PNNs in the UF, if any at all. The biological relevance of this finding is not yet understood, especially since NECTIN3 is very lowly expressed in OLs, and warrants further investigation.

MicroRNAs have emerged as important post-transcriptional epigenetic regulators of gene expression, and a large body of literature has linked differential levels of microRNA to early life adversity, depression and suicidality ^66–69^. In fact, it has been demonstrated that microRNAs can mediate the link between early life adversity and depression/suicidal behavior ^66^. MIR646HG, the host gene for MIR646, was downregulated in DS-CA in OPC, OL, and LAMP5+ inhibitory neurons in the current dataset. Via genome-wide association studies, variants in MIR646HG have been associated with depression ^70^, externalizing behaviors ^71^, cognitive abilities ^72^, and educational attainment ^73^. A computationally predicted target of MIR646 is NECTIN3. Despite their canonical role in gene silencing, microRNAs and their target genes can have other relationships ^74^. For example, they can be engaged in a double-negative feedback loop in which the microRNA and its target can reciprocally regulate each other’s expression ^75^. Other mechanisms, such as mRNAs competing for a common microRNA binding site, might be relevant ^76^. Importantly, we do not know how much of the mature microRNA is present, which is critical as it is the mature form that regulates expression ^67^. Moreover, without knowing whether the expression of the MIR646HG correlates well with the target gene’s protein level, our interpretation is limited. Furthermore, since we measured nuclear RNA, and microRNA silencing mostly occurs in the cytoplasm ^77^, the expression of the target gene in this dataset may not be representative of the cytoplasmic expression nor of the protein levels of this gene. Therefore, more experimental evidence is needed to firmly link MIR646 to NECTIN3 expression and to understand the regulatory mechanism at play.

Despite these interesting DEGs, our UF histology, gene expression, and ultrastructure findings for CA/depression are in contrast to what has been previously observed in cortical WM. Long-lasting myelin-related effects of CA have been observed in ventromedial prefrontal cortex WM and anterior cingulate cortex (ACC) WM ^22,78,79^. In the human ACC WM, a history of severe CA was associated with long-lasting changes including a decrease in SOX10+ OL density, thinner myelin and higher g-ratio in small caliber fibers, along with changes to OL-lineage cell methylation and myelin-related gene expression in the ACC grey matter ^22^. Since myelin-related lasting cellular/molecular traces of CA were not detected in the UF, we can state that it appears to be affected differently from cortical WM.

We must therefore reconcile our findings with previous neuroimaging findings that show changes in fractional anisotropy associated with CA in the UF. Not all neuroimaging studies that investigate childhood maltreatment, or other early life stressors, report microstructure changes in the UF ^80–82^, and the directionality and laterality of changes reported are also inconsistent.

However, assuming the studies with positive results would indeed replicate, it is of importance to note that these studies have been mainly conducted in infants ^83,84^, children/adolescents ^4,5,10,13,14,85^, and young adults ^9,12^, while the subjects in the current study are often decades removed from the initial trauma. We thus speculate that the myelin-related UF neuroimaging findings of childhood maltreatment might represent transient, dynamic changes in maturation rate, rather than long-lasting neurobiological correlates that can be observed postmortem. Large scale, longitudinal neuroimaging studies that run well into middle/late adulthood would be able to better ascertain the temporal dynamics of the UF.

What biological mechanisms could underlie the differences observed in myelin-related CA changes between cortical WM and the UF? Human OL-lineage cells, microglia, and astrocytes from different CNS WM regions show different subtype composition and transcriptomic signatures ^86,87^, which affects the ways in which they communicate with each other via signaling molecules ^88^. It is plausible that regional heterogeneity in the expression of genes that trigger the switch to activity-dependent myelination (e.g., NRG1, ERBB receptors, NMDA receptor subunits) may play a role ^89–91^. Other microenvironmental differences, including the proximity to vasculature may be especially relevant given that the expression of the endothelin peptide, expressed by CNS vascular cells, is coupled to neuronal activity via increased blow flow, which in turn regulates the number of sheaths extended per OL ^92^. How regional OL heterogeneity might alter the short-and long-term response of these cells to the effects of CA should be the focus of future investigations.

In contrast to the narrower effect of group, we found prominent effects of age across all experimental modalities, with an emphasis on myelin-related alterations. It has been shown that the effects of aging are most pronounced in anterior regions of the brain as compared to posterior regions, and that WM tracts are particularly vulnerable to the effects of aging when compared to grey matter ^26,27^. In fact, recent work has shown that WM tracts have a faster aging velocity compared to surrounding grey matter, with respect to a common aging score (i.e., a brain-wide aging gene “fingerprint”), which was predominant in glia ^27^. Furthermore, we identified aging DEGs only in glia, and it has previously been established that glia are more impacted by age than neurons and/or that they are affected earlier than neurons (Salas et al., 2020; Soreq et al., 2017). As such, it is not surprising that the UF would demonstrate marked molecular changes across the age span.

We found a significant positive relationship between NogoA+ OL density and age in our histology results, indicating OL accumulation with age, which has been previously documented ^26,93–98^. Given that we observed decreases in the expression of genes coding for myelin-constituent proteins, it is plausible that more OLs are needed to maintain myelin integrity. In other words, perhaps OLs accumulate with age as individual OLs become less “efficient” at myelination.

Moreover, we report that that axon diameter and g-ratio both decrease across the age span. Studies have shown fiber density in the optic nerve of macaque monkeys was reduced in old compared to young subjects (Sandell & Peters, 2000) and similarly fiber density in the corpus callosum and cingulum bundle decreased with age (Bowley et al., 2011). This decreased fiber density is also observed in human aging (Fan et al., 2020). Some studies show that thicker fibres are maintained at the cost of losing thinner fibres (Marner et al., 2003, Stahon et al., 2016, Fan et al. 2020). In contrast, in the UF we observe a shift towards smaller axons overall, with a potential loss of larger axons, though we cannot detect which axons have been lost over time due to the cross-sectional nature of this work. Of note, we imaged the terminations of the temporal segment of the UF, where fibers tend to fan out, which may not be representative of the entire length of the fiber. Nonetheless, we speculate that the decline in GJC2 (connexin 47) with age observed in the OL DEGs might in part mediate the observed shift in UF axon diameters with age. Inhibiting connexin 47 on OLs can impede the flow of nutrients to axons in aglycemic conditions *in vitro* ^99^. Furthermore, connexin 47 is frequently coupled with connexin 43 on astrocytes, forming a heterotypic “pan-glial” network, known to be important in maintaining the supply of energy to axons ^100,101^. Therefore, a decrease in inter-glial signalling and metabolic transport may be associated with shifts in UF axon ultrastructure.

Furthermore, there is strong evidence that paranodal reorganization occurs with aging ^102,103^, and several of the aging DEGs resolved in the current study would support this notion. Aging DEGs include CNTN1, a central adhesion molecule in the tripartite paranodal adhesion complex, NKX6-2, the transcription factor required for normal paranodal structure ^104^, and other components known to be enriched at the juxtaparadonal region (e.g., EPB41L2, HAPLN2) ^105^.

While axon guidance molecules are known to have a role in synapse maintenance in the adult brain ^106–108^, we suspect that these molecules might also contribute to the remodelling of the axo-myelin paranode. For example, DCC signalling is necessary for the maintenance of paranodal adhesion ^109^. Therefore, paranodal remodelling may be occurring in the UF alongside the numerous other myelin-associated aging processes.

Interestingly, we have previously observed alterations in levels of myelin phospholipid fatty acids with age in the UF ^53^, specifically an increase in monounsaturated fatty acids (e.g., C16:1n-7, C18:1n-7), and a decrease in long-chain omega-6 polyunsaturated fatty acids (e.g. C 22:4n-6). In our age DEGs, we identified several lipid transport/remodeling genes (including CERS1, HACD1, STARD4, ABCA9, PLD5, ATP8A2, DGKE). As such, changes to lipid homeostasis may be a particularly important part of UF myelin aging. However, we can probably attribute the decrease in long chain omega-6 fatty acids to peripheral ELOVL2 levels. The downregulation of ELOVL2 expression is one of the major markers of biological aging, established in both humans and rodents ^110^. In fact, in a study of the “methylation clock” framework of human epigenetic aging, it was found that four of the top ten methylation markers were in the ELOVL2 regulatory element ^110^. As such, with age, methylation of ELOVL2 increases and expression decreases throughout the body, prominently in the liver ^110^. Since polyunsaturated fatty acids (PUFA) in the brain are mostly derived from circulation, it follows that a decrease in PUFA production in the liver would be reflected in the brain ^111^.

By integrating the aging results of the current paper with those of Perlman et al. 2025 ^53^, we depict a summary schematic of UF aging changes in Figure 6, which encompasses changes at the level of RNA, lipids, proteins, cell density, and ultrastructure. It is unclear when these changes occur in the temporal sequence of aging or their place in the biological cascade of the aging brain. It is likely that changes in UF biophysical properties (e.g., protein-lipid interactions, lipid-lipid interactions, axon diameter/g-ratio) would impact the transmission of information across brain regions and may therefore, at least in part, underlie the cognitive deficits that occur in the aged brain ^112,113^. Further research on the genetic and epigenetic regulation of aging in the human UF would yield important insights into the coordination of these cellular, molecular, and ultrastructural shifts across the lifespan.

**Figure 6.**
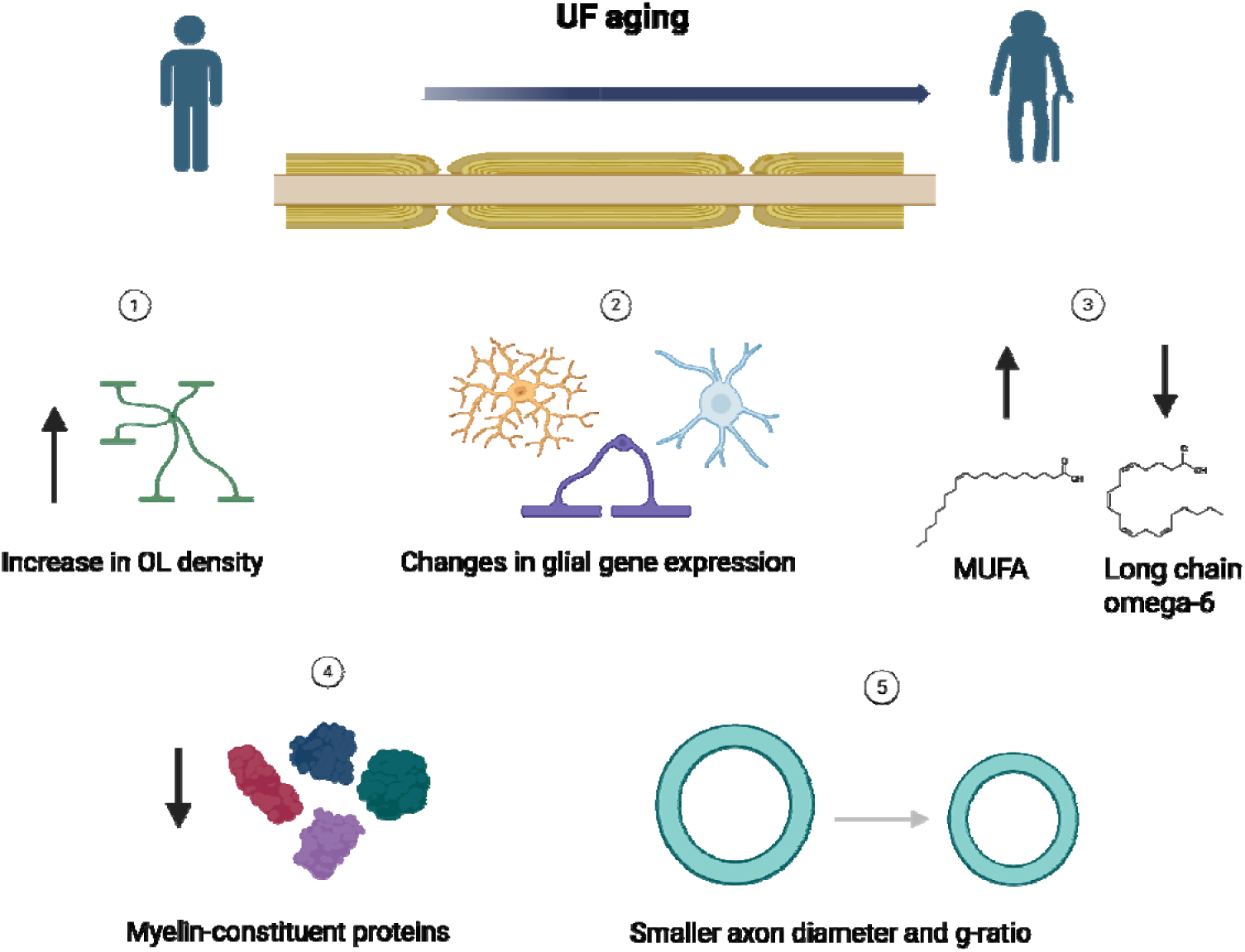
Summary schematic of age-related changes in the human postmortem UF. Diagram created in https://BioRender.com. Across all the UF experimental modalities, including those reported in our companion paper ^53^, throughout the age span we observe: 1) an increase in NogoA+ OL density; 2) changes in gene expression across glial (primarily OL-lineage) cell types that point to transcriptome changes in a wide variety of biological processes including ion channel transport, oxidative phosphorylation, ubiquitination, unfolded protein stress response, immune response, among others; 3) lipid changes with an overall pattern of an increase in monounsaturated fatty acids (MUFA) such as C16:1n-7 and C18:1n-7, and decreases in long chain omega-6 fatty acids such as C22:4n-6; 4) trends of decreasing myelin-constituent proteins and the genes that encode them; and 5) ultrastructural changes of UF myelin in which smaller diameter axons predominate and g-ratio decreases (myelin thickness shows decreasing relationship with age as well, but does not reach statistical significance). Altogether, this pattern of cellular, molecular, and ultrastructural findings suggests profound age-related alterations in the human UF.

This study is not without limitations. Due to limited tissue availability, we did not have the same cohort of subjects for all experimental approaches and parameters examined, preventing intra-subject comparisons and correlations. Furthermore, the maximum age in snRNAseq subject group (66 years) was far less than the maximum age in the stereology (90 years) and sf-CARS (85 years) cohorts, so we were likely not capturing the most extreme aging changes in gene expression. Furthermore, in the snRNAseq cohort, we only compared DS-CA and CTRL, thus preventing the attribution of DEGs specifically to effects of CA or of depression. Importantly, we were limited by the small sample size and may have been unable to pick up findings with small effect sizes. Another power issue might arise from the fact that we detected fewer axons per subject on average (<1000 axons per subject) as compared to a previous paper on the ACC ^22^ (∼2500-3000 axons/subject). Furthermore, we only sampled the left UF in these experiments, so we cannot immediately extend any conclusions to the right UF. Similarly, all conclusions are limited to the temporal segment of the UF, and as such cannot be extrapolated to the frontal or insular segments. This point is particularly relevant since a DTI study of early life stress sensitivity found effects in the frontal UF, but not the insular or temporal segments ^13^. Lastly, consistent with the known higher rates of suicide in males as compared to females ^114^, we had fewer female subjects in all cohorts. In the stereology cohort, we had no females. As such, it is important to replicate these studies in cohorts with more female subjects to better understand possible sex differences in this tract.

In summary, this study provides the first comprehensive cellular and molecular characterization of the human UF. We identified several between group differences by snRNAseq that will serve as interesting targets for follow-up, namely NECTIN3 and MIR646HG. However, we did not identify any group changes in myelin-related gene expression, nor in OL-lineage histology or ultrastructure. This absence of myelination-related changes indicates that the UF is affected differently from cortical WM. In contrast, we observed prominent myelin-age associations across all experimental modalities, adding new information to our understanding of the aging human brain. We believe that this study, together with our recent complementary work on the UF ^53^ will serve a foundational resource for future investigations of the human UF in both physiological and pathological conditions.

## Data availability

The raw and processed snRNAseq data is available as GSE319830 on the GEO repository.

## Code availability

Code used for the snRNAseq bioinformatic analysis is available at https://github.com/MGSSdouglas/Uncinate-Fasciculus-snRNAseq

The full model for the AxonDeepSeg segmentation model is available here: https://github.com/axondeepseg/model-seg-human-brain-cars.

## Acknowledgments

We would like to thank Sara-Maude Gagnon for her assistance in sf-CARS imaging. We would like to extend our thanks to the Douglas Bell-Canada Brain Bank staff, as well as the Molecular and Cellular Microscopy Platform (MCMP) at the Douglas Research Center. Most importantly, we thank the brain donors and their families for their most valuable gift.

## Author contributions

KP: conceptualization, methodology, formal analysis, investigation, project administration, writing-original draft; SBB: conceptualization, methodology, investigation, writing-review and editing; JK: methodology, investigation, writing-review and editing; JM: investigation, data curation, writing – review and editing; AC: conceptualization, methodology, software, validation, formal analysis, writing – review and editing; VPN: conceptualization, methodology, validation, investigation, writing – review and editing; MM: methodology, formal analysis, writing-review and editing; MM: methodology, software, validation; SJ: methodology; MAD: methodology, writing – review and editing; GF: methodology, writing-review and editing; AC: methodology, writing-review and editing; DV: methodology, investigation; JCA: project administration, methodology, software; DC: project administration, resources, funding acquisition, writing review and editing; RPB: Methodology, resources, writing-review and editing; CN: methodology, resources, writing-review and editing; GT: resources, funding acquisition, writing-review and editing; NM: conceptualization, project administration, resources, funding acquisition, writing-review and editing, supervision.

