## Supplementary Material for "A multimodal characterization of the human uncinate fasciculus"

**Supplementary Tables**

Supplementary Table 1 – Subject information for immunohistochemistry/stereology experiments. Data is represented as mean ± standard error of the mean. P-values are derived from one-way ANOVAs.

|  | **CTRL** | **DS** | **DS-CA** |
| --- | --- | --- | --- |
| **N total** | 13 | 13 | 16 |
| **Male/Female** | 13/0 | 13/0 | 16/0 |
| **Age** (p = 0.025) | 37.23 ± 4.04 | 54.00 ± 4.82 | 37.38 ± 2.69 |
| **pH** (p = 0.69) | 6.56 ± 0.068 | 6.56 ± 0.100 | 6.47 ± 0.095 |
| **PMI (h)** (p = 0.073) | 31.19 ± 6.32 | 49.64 ± 8.00 | 31.89 ± 3.81 |
| **N antidepressants** | 0 | 4 | 3 |

Supplementary Table 2- Antibody information for immunohistochemistry experiments.

| **Protein target** | **Manufacturer** | **Catalog #** | **Species** | **Concentration** |
| --- | --- | --- | --- | --- |
| **PDGFRα** | R&D Systems | AF-307-NA | Goat | 1 in 100 |
| **NogoA** | Millipore | ab5888 | Rabbit | 1 in 500 |

Supplementary Table 3 - Subject information for snRNAseq. Data is represented as mean ± standard error of the mean. P-values are derived from Welch’s two sample t-test.

|  | **CTRL** | **DS-CA** |
| --- | --- | --- |
| **N total** | 16 | 19 |
| **Male/Female** | 5/11 | 7/12 |
| **Age** (p = 0.94) | 47.4 ± 2.99 | 47.1 ± 2.43 |
| **pH** (p = 0.68) | 6.38 ± 0.081 | 6.44 ± 0.099 |
| **PMI (h)** (p =0.27) | 54.2 ± 5.58 | 62.4 ± 4.80 |
| **N antidepressants** | 0 | 10 |

Supplementary Table 4 – Subject information for sf-CARS experiment. Data is represented as mean ± standard error of the mean. P-values are derived from one-way ANOVAs.

|  | **CTRL** | **DS** | **DS-CA** |
| --- | --- | --- | --- |
| **N total** | 6 | 13 | 11 |
| **Male/Female** | 3/3 | 10/3 | 4/7 |
| **Age** (p = 0.28) | 61.2 ± 7.27 | 56.8 ± 3.51 | 50.2 ± 3.93 |
| **pH** (p=0.58) | 6.49 ± 0.14 | 6.42 ± 0.056 | 6.56 ± 0.13 |
| **PMI (h)** (p=0.73) | 63.4 ± 8.25 | 62.0 ± 5.97 | 69.2 ± 7.55 |
| **N antidepressants** | 0 | 6 | 5 |

Supplementary Table 5 – Sequencing statistics by batch/pool from cellranger v8.0.1 and cellbender v0.3.

| **Test** | **n nuclei (cellranger)** | **reads/nuclei (cellranger)** | **genes/nuclei (cellranger)** | **sequencing saturation (%) (cellranger)** | **n nuclei (cellbender)** | **ambient RNA % (cellbender)** |
| --- | --- | --- | --- | --- | --- | --- |
| **UF pool 1** | 12783 | 79045 | 2718 | 71.1 | 19484 | 4.4 |
| **UF pool 2** | 6552 | 149875 | 2315 | 84.3 | 9219 | 4.14 |
| **UF pool 3** | 26046 | 79803 | 2488 | 79.5 | 28026 | 12.04 |
| **UF pool 4** | 11787 | 190576 | 2446 | 86.9 | 13450 | 3.3 |
| **UF pool 5** | 13963 | 211323 | 2774 | 84.5 | 17881 | 4.71 |
| **UF pool 6B** | 7584 | 150096 | 1922 | 89.6 | 8730 | 4.65 |
| **UF pool 7A** | 10260 | 167991 | 2019 | 89.8 | 15452 | 3.1 |
| **UF pool 8A** | 3262 | 576563 | 1883 | 97 | 9652 | 2.12 |
| **UF pool 9A** | 9953 | 216416 | 1939 | 92.4 | 14578 | 2.15 |
| **UF pool 10B** | 9257 | 211299 | 2120 | 91.7 | 14128 | 3.32 |
| **UF pool 11A** | 7299 | 259830 | 2123 | 92.8 | 10168 | 2.99 |
| **UF pool 12B** | 5551 | 379681 | 2003 | 94.9 | 10654 | 2.23 |

Supplementary Table 6 – Number of nuclei per individual cluster listed in descending frequency.

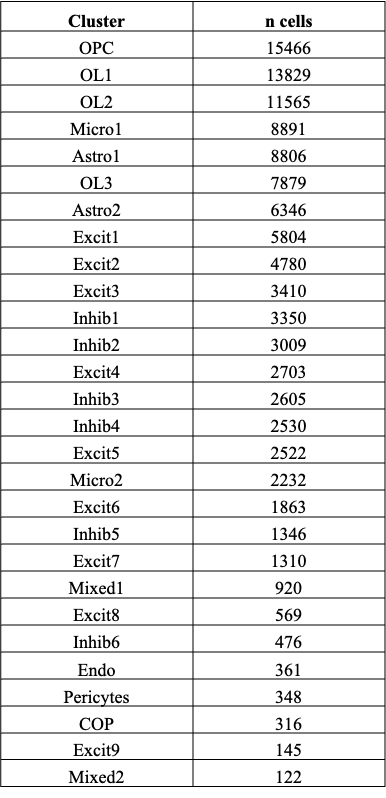

Supplementary Table 7 – Number of nuclei per broad cell type cluster.

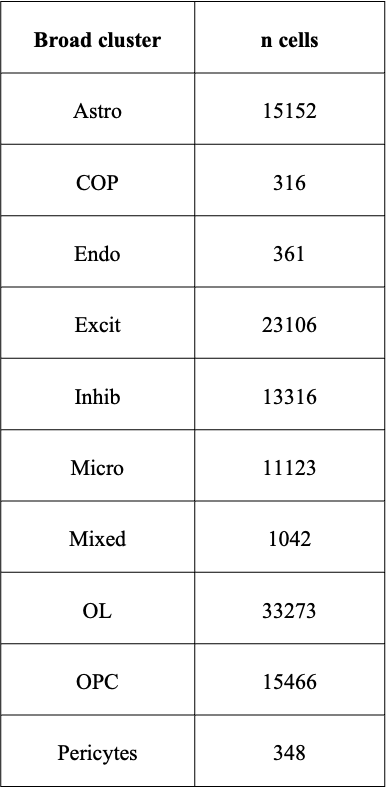

Supplementary Table 8 – Table showing individual neuron clusters and the name of their most concordant neuronal cluster from the MTG SEA-AD dataset mapping by Jaccard similarity index. The number of nuclei for each annotation is listed.

| Cluster name | Number of nuclei per cluster | Corresponding MTG-SEA-AD cluster name | Number of nuclei mapped to MTG SEA-AD cluster |
| --- | --- | --- | --- |
| Excit1 | 5804 | L6 IT | 3706 |
| Excit2 | 4780 | L6 CT | 2489 |
| Excit3 | 3410 | L2/3 IT | 4593 |
| Excit4 | 2703 | L5 IT | 3701 |
| Excit5 | 2522 | L6 IT Car3 | 2607 |
| Excit6 | 1863 | L4 IT | 1929 |
| Excit7 | 1310 | L6b | 3795 |
| Excit8 | 569 | L5/6 NP | 564 |
| Excit9 | 145 | L5 ET | 228 |
| Inhib1 | 3350 | Vip | 2980 |
| Inhib2 | 3009 | Pvalb | 2897 |
| Inhib3 | 2605 | Lamp5 | 2119 |
| Inhib4 | 2530 | SST | 2553 |
| Inhib5 | 1346 | Lamp5 lhx6 | 1277 |
| Inhib6 | 476 | Chandelier | 488 |

Supplementary Table 9 – Table showing DEGs between DS-CA and CTRL for individual clusters. This table shows the gene name, the cluster name, the locally adjusted p-value, and the logFC. Genes are ordered by ascending adjusted p-value. AS = antisense, DT = divergent transcript.

| gene | cluster_id | adj.P.Val | logFC |
| --- | --- | --- | --- |
| PCLO | OL3 | 0.00432022 | -1.2989789 |
| ARHGAP26 | OL3 | 0.00432022 | -1.2897093 |
| NECTIN3 | OL3 | 0.00432022 | -0.7865238 |
| PSRC1 | OL3 | 0.00486357 | 0.79536096 |
| LINC00844 | OL3 | 0.00486357 | 0.99942737 |
| CPB2-AS1 | OL1 | 0.00658671 | 0.50546217 |
| NOX4 | OL3 | 0.00712705 | -1.1625801 |
| TMEM178A | OL3 | 0.00737658 | -1.1230657 |
| DYNLL1 | Inhib3 | 0.00755768 | 0.80050632 |
| DENND2A | OL3 | 0.00838266 | 0.6864863 |
| HIVEP1 | OL3 | 0.01205577 | 0.96258394 |
| RPL30 | Inhib3 | 0.01867669 | 0.88004729 |
| RPS12 | Inhib3 | 0.01867669 | 1.00257772 |
| TOMM7 | Inhib3 | 0.01867669 | 1.09711169 |
| HSPA1B | OL3 | 0.01955824 | -1.8439451 |
| MIR646HG | Inhib3 | 0.03998279 | -0.9178635 |
| PRPF8 | Inhib3 | 0.03998279 | 0.67688424 |
| PPA1 | Inhib3 | 0.03998279 | 0.68805552 |
| NECTIN3 | OL1 | 0.04652743 | -0.7576972 |
| SRP14 | OPC | 0.04670915 | 0.88894588 |
| LINC01322 | Inhib5 | 0.04979836 | -0.8247817 |
| ALCAM | Inhib5 | 0.04979836 | -0.6750883 |
| KCND2 | Inhib5 | 0.04979836 | -0.457289 |
| PLCG2 | Inhib5 | 0.04979836 | 1.67548418 |
| RPL26 | Inhib3 | 0.05454511 | 1.09885856 |
| KIT | Inhib1 | 0.05970845 | -0.6950833 |
| USP6NL | OL3 | 0.06048248 | -0.8694783 |
| CDH2 | Inhib3 | 0.06593259 | -0.4267676 |
| LRIG2 | Inhib3 | 0.06593259 | 0.70563267 |
| RPS28 | Inhib2 | 0.07467633 | 1.16451913 |
| GABPB1-AS1 | Inhib2 | 0.07467633 | 1.43923608 |
| ENSG00000284977 | OL3 | 0.07845417 | -2.1130737 |
| MIR646HG | OL3 | 0.07845417 | -1.3005977 |
| ENSG00000289605 | OL3 | 0.07845417 | -1.0841647 |
| RELL1 | OL3 | 0.07845417 | -0.942812 |
| TRIM36 | OL3 | 0.07845417 | -0.7788669 |
| VOPP1 | OL3 | 0.07845417 | -0.6972769 |
| COPB2-DT | OL3 | 0.07845417 | 0.40793824 |
| ENSG00000268518 | OL3 | 0.07845417 | 0.71276365 |
| AMZ1 | OL3 | 0.07845417 | 1.06715533 |
| TNIK | Inhib3 | 0.07858045 | -0.7808887 |
| RPS7 | Inhib3 | 0.07858045 | 0.69558916 |
| MIR646HG | OL1 | 0.0862345 | -1.0096671 |
| STARD13 | OL3 | 0.0870871 | 0.3523524 |
| TOM1L2 | Inhib5 | 0.09297864 | 0.68322801 |
| LINC00486 | Inhib5 | 0.09297864 | 0.86012939 |
| FSTL5 | OL3 | 0.09749169 | -1.4342811 |
| LINC02511 | OL3 | 0.09749169 | -0.9108268 |
| PHGDH | OL3 | 0.09749169 | -0.6275135 |

Supplementary Table 10 – Table showing DEGs between DS-CA and CTRL for broad clusters. This table shows the gene name, the cluster name, the locally adjusted p-value, and the logFC. Genes are ordered by ascending adjusted p-value. AS = antisense.

| Gene | Cluster | adj.P.Val | logFC |
| --- | --- | --- | --- |
| NECTIN3 | OL | 0.00436505 | -0.7710088 |
| SRP14 | OPC | 0.03224629 | 0.90027851 |
| FSTL5 | OL | 0.0449736 | -1.660986 |
| MIR646HG | OL | 0.0449736 | -1.1051 |
| RELL1 | OL | 0.05257687 | -1.1154637 |
| ENSG00000289605 | OL | 0.05257687 | -1.4062444 |
| C1orf174 | OL | 0.05257687 | 0.88152387 |
| EPS8 | OL | 0.05257687 | -0.5532322 |
| ENSG00000268518 | OL | 0.05257687 | 0.69507598 |
| SAMD4A | OL | 0.05886557 | -0.6872799 |
| CEP164 | OL | 0.06404538 | 0.46284554 |
| GRIN2A | OPC | 0.06760199 | 1.04054693 |
| SRSF2 | OPC | 0.06760199 | 0.5236616 |
| DSEL-AS1 | OPC | 0.06760199 | -0.5089832 |
| NDUFA13 | OPC | 0.06760199 | 0.67053315 |
| MIR646HG | OPC | 0.06760199 | -1.2230183 |
| RNF152 | OL | 0.06965061 | -0.7025029 |
| ENSG00000248458 | OPC | 0.08616575 | 0.96998453 |
| GNPDA2 | OPC | 0.08616575 | -0.4164708 |
| ENSG00000228541 | OL | 0.09874493 | -0.6308906 |

Supplementary Table 11 – Table showing significant age DEGs for individual clusters. This table shows the gene name, the cluster name, the locally adjusted p-value, and the logFC. Genes are ordered by ascending adjusted p-value. AS = antisense, DT = divergent transcript.

| gene | cluster_id | adj.P.Val | logFC |
| --- | --- | --- | --- |
| IGFBP7 | Astro1 | 0.00334869 | 0.57631942 |
| ADAMTS9-AS2 | OPC | 0.00438955 | 0.34202438 |
| GPC5 | OL1 | 0.00486111 | 1.07320751 |
| COL16A1 | OL1 | 0.00486111 | -0.5101193 |
| ARHGEF3 | OL1 | 0.00593103 | -0.8705412 |
| RORB | OL1 | 0.00593103 | 1.44841842 |
| STON1 | OL1 | 0.0067227 | 0.67661771 |
| MACROH2A2 | OL1 | 0.0072494 | 0.93905097 |
| SDK1 | OL1 | 0.00862595 | 1.01759711 |
| FXYD6 | OL1 | 0.00862595 | 1.01753968 |
| VASH1-DT | OPC | 0.01349104 | -0.3361658 |
| SFRP1 | OL1 | 0.01696319 | 0.61093659 |
| RELN | OL1 | 0.01696319 | 0.63540015 |
| PARD3 | OL1 | 0.01724784 | 1.03016971 |
| SPATA6 | OL1 | 0.01791646 | -0.3426437 |
| ATP5ME | OL1 | 0.01791646 | -0.4577292 |
| LINC02821 | OL1 | 0.01791646 | -0.5945487 |
| GDNF-AS1 | OL1 | 0.01791646 | 0.56418286 |
| HEPN1 | OL1 | 0.01791646 | 0.48744075 |
| EPB41L2 | OL1 | 0.01791646 | -0.2610451 |
| GUCY1A2 | OL1 | 0.01835921 | 0.90508218 |
| SLC6A15 | OL1 | 0.01906879 | 0.9585377 |
| PCAT1 | Micro1 | 0.02079458 | -1.1571856 |
| CNTN1 | OL1 | 0.02459414 | 0.78721577 |
| PAK5 | OL1 | 0.02459414 | 0.99488605 |
| HACD1 | OL1 | 0.02459414 | 0.47539269 |
| GNG7 | OL1 | 0.02472766 | -0.321743 |
| MT-CYB | Astro2 | 0.02578929 | 1.23967408 |
| MYRIP | OL1 | 0.02716245 | 0.59185987 |
| ART3 | OL1 | 0.03054554 | 0.64761476 |
| LZTFL1 | OL1 | 0.03263737 | 0.55986463 |
| LINC03021 | OL1 | 0.03263737 | 1.42024204 |
| ENSG00000264513 | OL1 | 0.03263737 | 0.48571588 |
| UBE3B | OL1 | 0.0337581 | 0.32027368 |
| TNIK | OL1 | 0.0341905 | 0.41687187 |
| CNKSR3 | OL1 | 0.04150433 | -0.3067773 |
| MBP | OL1 | 0.04337893 | -0.2112056 |
| PTPRM | OL1 | 0.04337893 | 0.9825254 |
| GAS7 | OL1 | 0.04507732 | -0.2717252 |
| GSN | OL1 | 0.04507732 | -0.2368667 |
| FLNB | OL1 | 0.04537344 | 0.35281435 |
| COL10A1 | OL1 | 0.04564921 | 0.67346533 |
| LMCD1-AS1 | OL1 | 0.04564921 | -0.2550002 |
| STARD4 | OL1 | 0.04564921 | 0.80554882 |
| LRCH2 | OL1 | 0.04564921 | 0.3671538 |
| CABCOCO1 | OL1 | 0.04564921 | 0.59996212 |
| CRIM1 | OL1 | 0.04564921 | 0.39561804 |
| MYO16 | OPC | 0.04734561 | 0.23695013 |
| FEZ1 | OL1 | 0.04807437 | -0.2456553 |
| TGFBR2 | OL1 | 0.04807437 | 0.54883342 |
| PDZRN4 | OL1 | 0.0507665 | 0.92283447 |
| ENSG00000285367 | OPC | 0.05127933 | 0.62437229 |
| DNAJB2 | OL1 | 0.05234478 | -0.2902325 |
| DCC | OL1 | 0.05385162 | 0.86324516 |
| H2BC4 | OL1 | 0.054079 | 0.40459002 |
| DTNA | OL1 | 0.05504634 | -0.4442852 |
| C1GALT1 | OL1 | 0.05682772 | -0.2249134 |
| KCNAB1 | OL1 | 0.05688847 | 0.47473161 |
| INTS12 | OL1 | 0.05688847 | 0.41625199 |
| TF | OL1 | 0.05709842 | -0.2105821 |
| PAQR3 | OL1 | 0.06358315 | 0.537703 |
| FMNL2 | OL1 | 0.06424243 | -0.1991701 |
| ENSG00000259678 | OL1 | 0.0654481 | -0.4514592 |
| LINC02609 | OL1 | 0.0654481 | 0.66622788 |
| ENSG00000251600 | OL1 | 0.06698663 | -0.4046335 |
| LINC00844 | OL1 | 0.06698663 | 0.38795461 |
| ENSG00000251652 | OL1 | 0.06746644 | -0.8851451 |
| MIR31HG | OL3 | 0.06892909 | 0.78058873 |
| SLC6A15 | OL3 | 0.06892909 | 0.91442983 |
| RORB | OL3 | 0.06892909 | 1.21046839 |
| EPB41L2 | OL3 | 0.06892909 | -0.2400342 |
| PLD5 | OPC | 0.07055566 | 0.33754921 |
| SPARCL1 | OPC | 0.07055566 | 0.25343403 |
| ENSG00000251600 | OPC | 0.07055566 | -0.3151331 |
| LINC00299 | OPC | 0.07055566 | 0.41884251 |
| MERTK | OPC | 0.07055566 | 0.4784102 |
| SLC24A4 | OPC | 0.07055566 | 0.60374173 |
| PCDH10-DT | OPC | 0.07055566 | -0.2864385 |
| FRMD6-AS2 | OPC | 0.07055566 | 0.39295567 |
| LINC01470 | OL1 | 0.07139326 | 0.542253 |
| ABCA9 | OL1 | 0.07166222 | 0.88434826 |
| DISP1 | OL1 | 0.07225217 | 0.51572697 |
| ENSG00000286446 | OL1 | 0.07225217 | 0.56478541 |
| SLC7A11 | OPC | 0.07248092 | 0.76652673 |
| THBS4 | OPC | 0.07248092 | 0.2624695 |
| DNM2 | OL1 | 0.07571076 | -0.2437192 |
| DGKE | OL1 | 0.07571076 | 0.57008305 |
| ARHGAP22 | OL1 | 0.07571076 | -0.2661316 |
| EYS | OL1 | 0.07571076 | 0.72231227 |
| MED4 | OL1 | 0.07712682 | 0.32033621 |
| ARL17A | OL1 | 0.07774577 | 0.58195421 |
| ENSG00000254604 | OL1 | 0.07774577 | 0.54501744 |
| ADAMTS17 | OL1 | 0.07774577 | 0.66075358 |
| CNTN3 | OL1 | 0.07774577 | 0.86823668 |
| DOCK3 | OL1 | 0.07911552 | -0.2162504 |
| ATP8A2 | OL1 | 0.08161565 | 0.89450841 |
| AUH | OL1 | 0.08194428 | 0.23209136 |
| PLXNB1 | OL1 | 0.08194428 | -0.2508712 |
| PTPRK-AS1 | OL1 | 0.08194428 | 0.59627184 |
| SSX2IP | OL1 | 0.08755216 | 0.54399626 |
| CTBS | OL1 | 0.08755216 | 0.59123017 |
| BTD | OL1 | 0.08755216 | 0.30131294 |
| TGDS | OL1 | 0.08755216 | 0.43777623 |
| RNF212 | OL1 | 0.08755216 | -0.7422091 |
| ENSG00000288891 | OL1 | 0.08755216 | 0.65063946 |
| ENSG00000287720 | OL1 | 0.08755216 | 0.61380053 |
| S100A6 | OL1 | 0.08868301 | 1.08772381 |
| SAMHD1 | OL1 | 0.08868301 | 0.64848191 |
| NINL | OL1 | 0.08985514 | 0.4709675 |
| CDH26 | OL1 | 0.09038759 | 0.39241746 |
| ZNF302 | OL1 | 0.09053733 | 0.29554405 |
| ZNF529-AS1 | OPC | 0.09428413 | 0.29805316 |
| ZNF676 | OPC | 0.09428413 | 0.9541429 |
| ENSG00000259124 | OPC | 0.09655764 | -0.3014546 |
| PON3 | OPC | 0.09655764 | 0.78566077 |
| SYTL3 | Micro1 | 0.09748255 | 0.6197264 |
| MIR34AHG | Micro1 | 0.09748255 | 0.46388797 |
| RRM2B | Micro1 | 0.09748255 | 0.47402936 |
| TTC7B | Micro1 | 0.09748255 | 0.35156318 |
| DAB2IP | Micro1 | 0.09748255 | 0.92103083 |
| PLOD2 | OL1 | 0.09762533 | 0.67497584 |
| TSBP1-AS1 | OL3 | 0.09873452 | -0.4086862 |

Supplementary Table 12 – Table showing significant age DEGs for broad clusters. This table shows the gene name, the cluster name, the locally adjusted p-value, and the logFC. Genes are ordered by ascending adjusted p-value. AS = antisense. DT = divergent transcript.

| gene | cluster_id | adj.P.Val | logFC |
| --- | --- | --- | --- |
| ADAMTS9-AS2 | OPC | 0.006681 | 0.3461926 |
| VASH1-DT | OPC | 0.01402998 | -0.3401865 |
| LINC02821 | OL | 0.01628883 | -0.5884005 |
| SLC6A15 | OL | 0.01628883 | 1.0168456 |
| FXYD6 | OL | 0.01628883 | 1.13603244 |
| GUCY1A2 | OL | 0.01628883 | 1.13814169 |
| NKX6-2 | OL | 0.01628883 | -0.3623572 |
| TMEM26 | OL | 0.01628883 | 1.50058926 |
| CRIM1 | OL | 0.01628883 | 0.42340426 |
| ARHGEF3 | OL | 0.01628883 | -0.5822268 |
| ENSG00000251600 | OPC | 0.01855553 | -0.325202 |
| MACROH2A2 | OL | 0.02371974 | 0.78916453 |
| KCNAB1 | OL | 0.02393254 | 0.64892683 |
| IGFBP7 | Astro | 0.02484107 | 0.54359606 |
| MYO16 | OPC | 0.02783402 | 0.24071102 |
| PLD5 | OPC | 0.02860499 | 0.34379984 |
| EPB41L2 | OL | 0.0310773 | -0.2887128 |
| RORB | OL | 0.0310773 | 1.48711326 |
| GDNF-AS1 | OL | 0.0310773 | 0.50914649 |
| LHPP | OL | 0.03254658 | -0.314306 |
| PPP1R14A | OL | 0.03356987 | -0.4171433 |
| LINC01792 | OL | 0.03356987 | 0.38573331 |
| CNKSR3 | OL | 0.03356987 | -0.2685868 |
| SFRP1 | OL | 0.03680276 | 0.56410579 |
| MEF2C-AS2 | OL | 0.03680276 | 0.75756792 |
| ENSG00000285367 | OPC | 0.03714199 | 0.61613759 |
| MAPT-AS1 | OL | 0.03910317 | 0.52825721 |
| KLHL13 | OL | 0.04178633 | 0.3928352 |
| STON1 | OL | 0.04178633 | 0.44814014 |
| HSP90B1 | OL | 0.04178633 | -0.2713825 |
| PCAT1 | Micro | 0.04653194 | -1.0929608 |
| MIR34AHG | Micro | 0.04653194 | 0.4867974 |
| ZNF423 | Micro | 0.04653194 | -1.1734815 |
| HAPLN2 | OL | 0.05628984 | -0.2913918 |
| RASSF2 | OL | 0.05704392 | -0.2526607 |
| MYH15 | OL | 0.05704392 | 0.41993986 |
| S100A6 | OL | 0.0590381 | 0.86804935 |
| SPP1 | OL | 0.0590381 | -0.3246974 |
| PAK5 | OL | 0.06002699 | 1.09358437 |
| XKR6 | OL | 0.06002699 | -0.2918314 |
| MICALL1 | OL | 0.06002699 | -0.329241 |
| DCC | OL | 0.06040824 | 0.71594757 |
| FEZ1 | OL | 0.06040824 | -0.2401885 |
| TMEM179 | OL | 0.06040824 | 0.91506889 |
| PDZRN4 | OL | 0.06040824 | 1.06347628 |
| KCND3 | OL | 0.06040824 | 0.92786576 |
| CERS1 | OL | 0.06040824 | -0.4409349 |
| NKD1 | OL | 0.06040824 | -0.3390777 |
| ATP5ME | OL | 0.06040824 | -0.3978606 |
| MARCHF3 | OL | 0.06040824 | 0.77625476 |
| EML5 | OL | 0.06040824 | 0.5085116 |
| LZTFL1 | OL | 0.06040824 | 0.38068819 |
| ART3 | OL | 0.06040824 | 0.48104614 |
| OR2L3 | OL | 0.06040824 | 0.68721713 |
| MTUS1 | OL | 0.06249108 | -0.2714251 |
| ATP8A2 | OL | 0.06249108 | 0.83540322 |
| ACTN2 | OL | 0.06361915 | -0.4053694 |
| SSBP3 | OL | 0.06361915 | -0.2632597 |
| ABHD17A | OL | 0.06639631 | -0.264221 |
| GJC2 | OL | 0.07075049 | -0.326817 |
| DBNDD2 | OL | 0.07075049 | -0.3110527 |
| COL28A1 | OL | 0.07075049 | 0.32967057 |
| SLC4A5 | OL | 0.07075049 | 0.2939005 |
| TNIK | OL | 0.07075049 | 0.42255906 |
| PRANCR | OL | 0.07075049 | 0.28561117 |
| EPHA3 | OL | 0.07075049 | 0.85485909 |
| KLRK1-AS1 | OL | 0.07075049 | 0.49953535 |
| CNTN1 | OL | 0.07277687 | 0.5933387 |
| MICALL2 | OL | 0.07277687 | 0.41903908 |
| LANCL1 | OL | 0.07277687 | -0.279664 |
| ROGDI | OL | 0.07277687 | -0.2945837 |
| SYTL3 | Micro | 0.07325709 | 0.60092023 |
| RRM2B | Micro | 0.07325709 | 0.45450087 |
| ADAMTS17 | Micro | 0.07325709 | 0.90776959 |
| SPARCL1 | OPC | 0.07474831 | 0.25359508 |
| LINC00299 | OPC | 0.07474831 | 0.42295782 |
| ENSG00000183308 | OL | 0.07507981 | 0.3829835 |
| DIRC3 | OL | 0.07775849 | 0.91937771 |
| OR2L5 | OL | 0.07825874 | 0.61626571 |
| STAT2 | OL | 0.07825874 | -0.2801809 |
| RSPO2 | OL | 0.07973345 | 0.57257729 |
| PCDH10-DT | OPC | 0.08075263 | -0.2874707 |
| ZNF676 | OPC | 0.08075263 | 0.96647745 |
| SOX13 | OL | 0.08429837 | -0.3744966 |
| MAP3K11 | OL | 0.08429837 | -0.26491 |
| TMOD1 | OL | 0.08429837 | 0.33394957 |
| SEPTIN7-DT | OL | 0.08741629 | 0.31016123 |
| PCCA-DT | OL | 0.08741629 | 0.42644974 |
| RAB40B | OL | 0.08879252 | -0.228103 |
| SLC24A4 | OPC | 0.08898452 | 0.62024971 |
| ZNF529-AS1 | OPC | 0.0896297 | 0.30412374 |
| SLC7A11 | OPC | 0.0896297 | 0.75879886 |
| FRMD6-AS2 | OPC | 0.0896297 | 0.39215977 |
| SLC1A3 | OPC | 0.0896297 | 0.22187343 |
| MERTK | OPC | 0.0896297 | 0.47443014 |
| TSBP1-AS1 | OL | 0.09013465 | -0.3578797 |
| SEMA6D | OL | 0.09088891 | 0.38589414 |
| MAPK8IP1 | OL | 0.09149995 | -0.2702069 |
| DGKE | OL | 0.09180987 | 0.47912021 |
| MDM4 | OL | 0.09274881 | -0.2141903 |
| TTYH1 | OL | 0.09274881 | -0.2501947 |
| ORC5 | OL | 0.09274881 | 0.20537232 |
| ENSG00000280206 | OL | 0.09274881 | -0.4904993 |
| ENSG00000264513 | OL | 0.09274881 | 0.36345581 |
| THBS4 | OPC | 0.09287343 | 0.25925058 |
| POU6F2 | OPC | 0.09287343 | 0.25702609 |
| NASP | OL | 0.09466266 | -0.242227 |
| GNG7 | OL | 0.09699524 | -0.2645416 |
| JARID2 | OL | 0.09699524 | -0.1863956 |
| MTA2 | OL | 0.09699524 | -0.3867084 |
| MBP | OL | 0.09768652 | -0.2968476 |
| XRRA1 | OL | 0.09879233 | 0.33681037 |
| MYL6B | OL | 0.09922418 | 0.65716833 |
| ENSG00000289410 | OL | 0.09922418 | 0.33875401 |
| LURAP1L-AS1 | OL | 0.09922418 | 1.43013633 |
| NACC2 | OL | 0.09994357 | -0.2813913 |
| PPFIA2 | OL | 0.09994357 | 0.2702962 |

Supplementary Table 13 – Overlap of significantly differentially expressed aging genes from the current study OL clusters (broad and individual) and from the oligodendrocyte (OLG) cluster from Ximerakis et al. (2019).

| **gene** | **cluster_id** | **logFC** | **adj.P.Val** | **logFC Ximerakis** | **p.adj Ximerakis** |
| --- | --- | --- | --- | --- | --- |
| ABHD17A | OL | 0.06639631 | -0.264221 | -0.060835 | 0.00123015 |
| ART3 | OL1 | 0.03054554 | 0.64761476 | 0.0124754 | 4.90051E-07 |
| ART3 | OL | 0.06040824 | 0.48104614 | 0.0124754 | 4.90051E-07 |
| CNKSR3 | OL1 | 0.04150433 | -0.3067773 | -0.0201417 | 1.31346E-07 |
| CNKSR3 | OL | 0.03356987 | -0.2685868 | -0.0201417 | 1.31346E-07 |
| CNTN1 | OL1 | 0.02459414 | 0.78721577 | -0.0494002 | 4.61455E-11 |
| CNTN1 | OL | 0.07277687 | 0.5933387 | -0.0494002 | 4.61455E-11 |
| DBNDD2 | OL | 0.07075049 | -0.3110527 | -0.0309446 | 0.01236479 |
| EML5 | OL | 0.06040824 | 0.5085116 | -0.0105897 | 0.08011004 |
| EPB41L2 | OL1 | 0.01791646 | -0.2610451 | 0.0334589 | 0.04872564 |
| EPB41L2 | OL3 | 0.06892909 | -0.2400342 | 0.0334589 | 0.04872564 |
| EPB41L2 | OL | 0.0310773 | -0.2887128 | 0.0334589 | 0.04872564 |
| FLNB | OL1 | 0.04537344 | 0.35281435 | 0.00115422 | 0.04188977 |
| FXYD6 | OL1 | 0.00862595 | 1.01753968 | -0.0096892 | 0.02654163 |
| FXYD6 | OL | 0.01628883 | 1.13603244 | -0.0096892 | 0.02654163 |
| GJC2 | OL | 0.07075049 | -0.326817 | -0.1450108 | 1.17734E-12 |
| GSN | OL1 | 0.04507732 | -0.2368667 | -0.4146582 | 1.45187E-97 |
| HACD1 | OL1 | 0.02459414 | 0.47539269 | -0.1376557 | 3.09924E-27 |
| HAPLN2 | OL | 0.05628984 | -0.2913918 | 0.18939913 | 8.05666E-20 |
| HSP90B1 | OL | 0.04178633 | -0.2713825 | 0.03534912 | 0.01058433 |
| KCNAB1 | OL1 | 0.05688847 | 0.47473161 | 0.02855307 | 5.60343E-08 |
| KCNAB1 | OL | 0.02393254 | 0.64892683 | 0.02855307 | 5.60343E-08 |
| LANCL1 | OL | 0.07277687 | -0.279664 | -0.0334619 | 0.00903193 |
| LZTFL1 | OL1 | 0.03263737 | 0.55986463 | 0.02092527 | 0.08568651 |
| LZTFL1 | OL | 0.06040824 | 0.38068819 | 0.02092527 | 0.08568651 |
| MAPK8IP1 | OL | 0.09149995 | -0.2702069 | -0.0393448 | 0.05493776 |
| MBP | OL1 | 0.04337893 | -0.2112056 | 0.10392351 | 9.03315E-52 |
| MBP | OL | 0.09768652 | -0.2968476 | 0.10392351 | 9.03315E-52 |
| MTA2 | OL | 0.09699524 | -0.3867084 | -0.0152257 | 0.04073422 |
| MYRIP | OL1 | 0.02716245 | 0.59185987 | 0.01191334 | 0.00192175 |
| NKD1 | OL | 0.06040824 | -0.3390777 | -0.0476661 | 0.04462927 |
| NKX6-2 | OL | 0.01628883 | -0.3623572 | -0.0586464 | 7.51484E-05 |
| PLOD2 | OL1 | 0.09762533 | 0.67497584 | -0.0020335 | 0.07581604 |
| PTPRM | OL1 | 0.04337893 | 0.9825254 | 0.00947041 | 0.03316033 |
| RASSF2 | OL | 0.05704392 | -0.2526607 | 0.04583734 | 0.01119415 |
| SEMA6D | OL | 0.09088891 | 0.38589414 | 0.07199019 | 2.2804E-05 |
| SSBP3 | OL | 0.06361915 | -0.2632597 | -0.0234397 | 0.00039722 |
| TNIK | OL1 | 0.0341905 | 0.41687187 | -0.0265997 | 0.00050832 |
| TNIK | OL | 0.07075049 | 0.42255906 | -0.0265997 | 0.00050832 |
| TTYH1 | OL | 0.09274881 | -0.2501947 | -0.1752341 | 2.7485E-47 |

**Supplementary Figures**

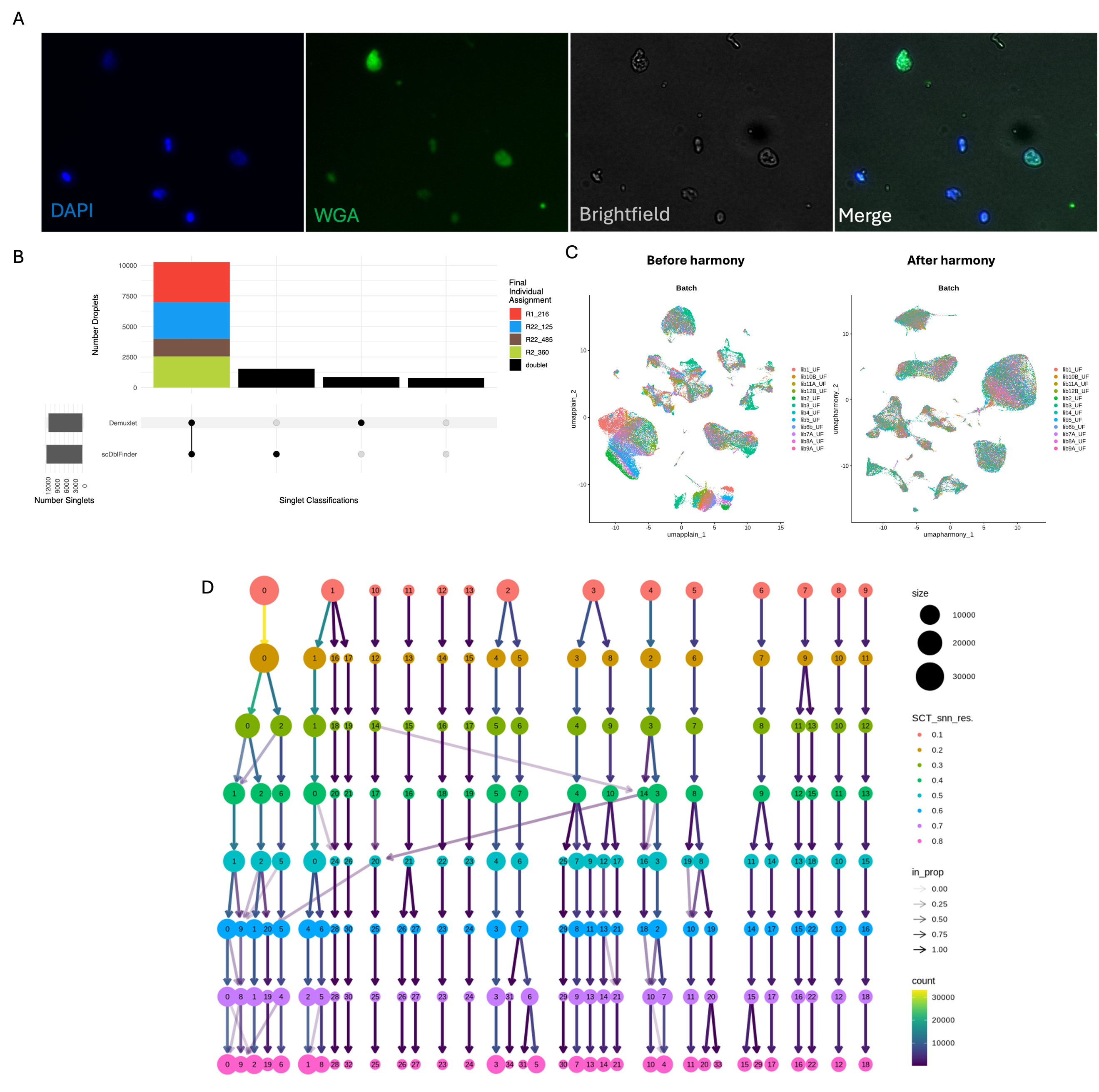

Supplementary Figure 1 – Quality control and preprocessing steps. A) Quality control on pool of dissociated nuclei. Micrographs at 20x magnification stained with DAPI (labels nuclei, blue), and wheat germ agglutinin (WGA, labels nuclear pore complexes, green). The suspension was also imaged in brightfield to visualize nuclei structure (grey). The merged image of fluorescence and brightfield micrographs is shown in the right-most panel. B) Example upset plot showing demultiplexing and dedoubletting results. The first bar on upset plot demonstrates singlets called by both tools (Demuxlet and scDblFinder), colored by subject ID (i.e., final individual assignment from Demuxlet). The second bar indicates singlets called by scDblFinder but not Demuxlet, the third bar indicates singlets called by Demuxlet but not scDblFinder, and the final bar indicates cells that were not deemed as singlets by either tool. The results between the two tools were combined using the “AnyDoublets” approach, indicating that a cell should be labelled as a doublet if either tool designates it a doublet. Doublet designations are colored in black. The horizontal bar plot on the far left illustrates the numbers of singlets called by each tool. C) UMAP plot colored by individual batch (library) before and after harmony integration. Each batch contains 4 subjects. D) Selecting optimal clustering resolution based on clustree plot stability. Each level of the tree corresponding to a clustering resolution level (from 0.1 at the top to 0.8 at the bottom). Each circle refers to a cluster, with the color of the circle representing the resolution of the clustering, and the size of the circle representing the number of nuclei (size) of each cluster. The opacity of the arrows represents the proportion of nuclei in one cluster that is moved to another cluster in the subsequent clustering resolution.

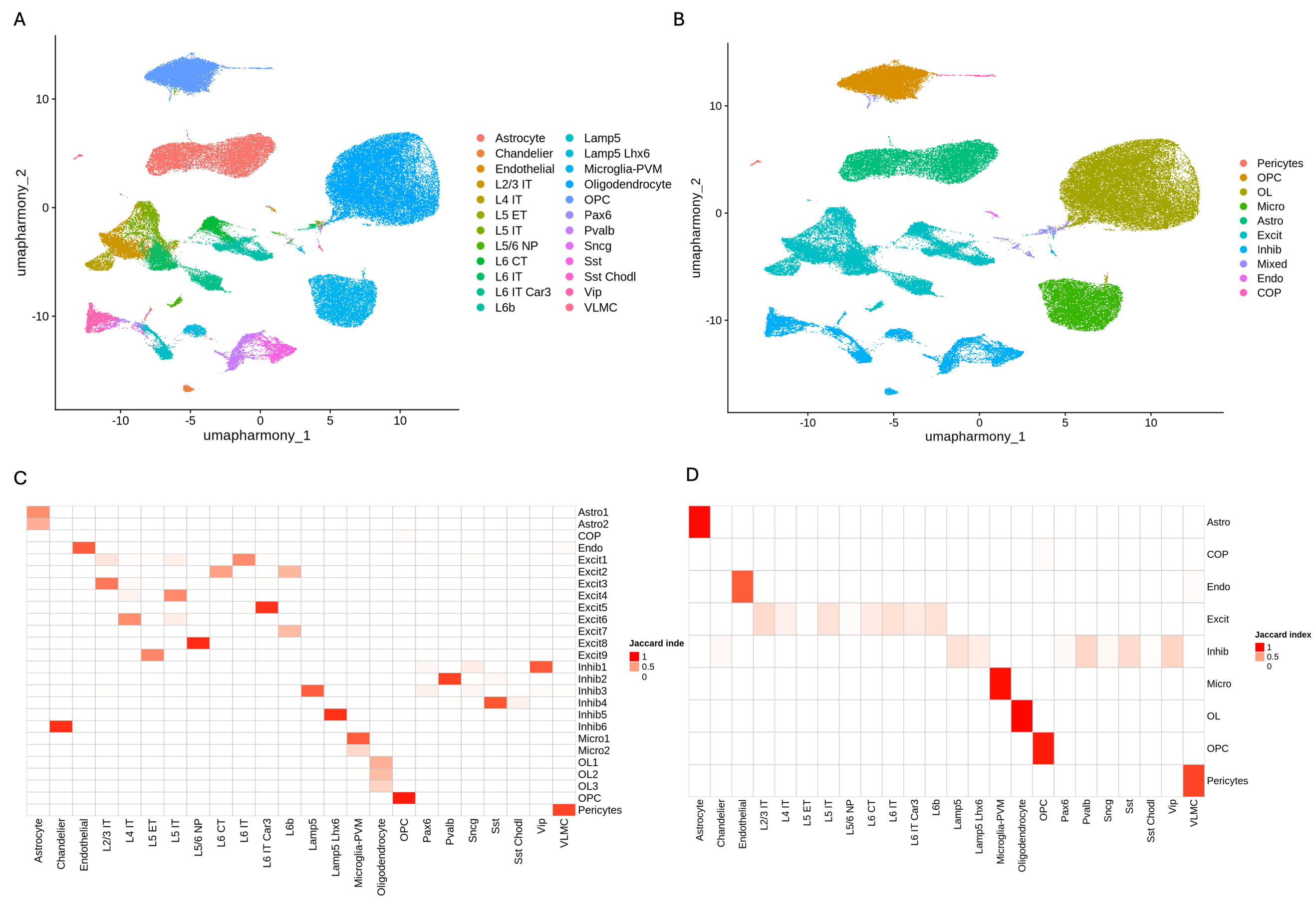

Supplementary Figure 2 – A) UMAP colored by cluster labels derived from the medial temporal gyrus (MTG) SEA-AD dataset. MapMyCells label transfer was performed with the 10X Human MTG SEA-AD dataset using the Deep Generative Mapping algorithm. B)

UMAP plot showing broad cell type clusters. Individual nuclei labelled by their broad cluster designations. C) Heatmap colored by Jaccard similarity index showing concordance between MTG SEA-AD and individual clusters. X-axis labels show the cluster labels from the MTG SEA-AD dataset mapped to the current dataset using MapMyCells. The y-axis labels refer to the individual cluster names from the current dataset. The cluster similarity is colored by the discrete Jaccard index values (0, 0.5, 1) with the scclusteval tool. The adjusted rand index between these two sets of labels is 0.615. D) Heatmap colored by Jaccard similarity index showing concordance between MTG SEA-AD and broad clusters. X-axis labels show the cluster labels from the MTG SEA-AD dataset mapped to the current dataset using MapMyCells. The y-axis labels refer to the broad cluster names from the current dataset. The cluster similarity is colored by the discrete Jaccard index values (0, 0.5, 1) with the scclusteval tool. The adjusted rand index between these two sets of labels is 0.821.

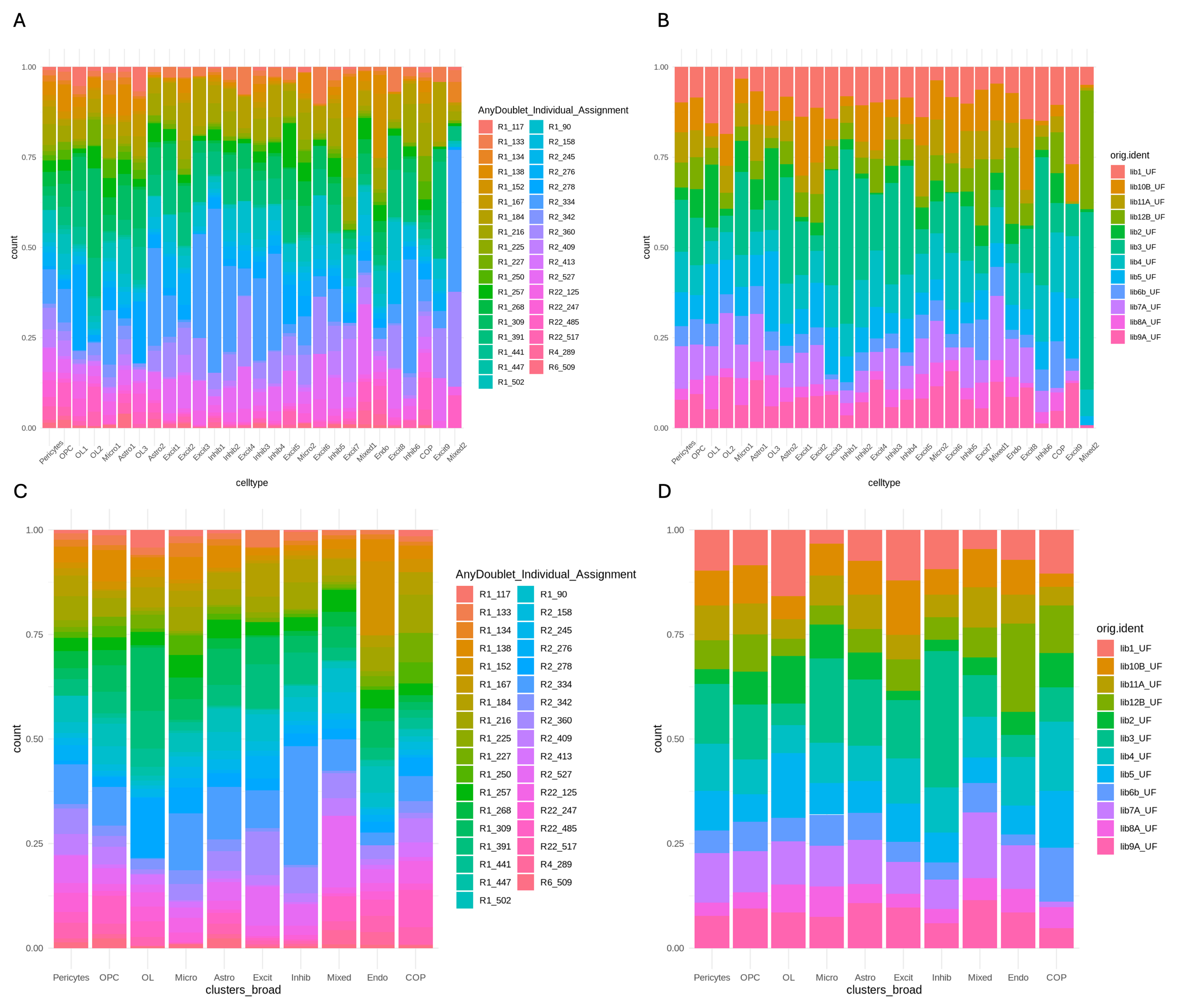

Supplementary Figure 3 – Distributions of subject and libraries in clusters. A) All individual clusters have contributions from all subjects, are not driven by a particular subject or batch, except for Mixed2 as shown by stacked bar plots showing distribution of subjects for individual cell type clusters. Each bar represents a cluster colored by the proportion of nuclei derived from each individual subject. B) Stacked bar plots showing distribution of batch (library) individual cell type clusters. Each bar represents a cluster colored by the proportion of nuclei derived from each individual batch. C) Broad clusters have contributions from all subjects and are not driven by a particular subject or batch as shown by stacked bar plots showing distribution of subjects for broad cell type clusters. Each bar represents a cluster colored by the proportion of nuclei derived from each individual subject. D) Stacked bar plots showing distribution of batch (library) broad cell type clusters. Each bar represents a cluster colored by the proportion of nuclei derived from each individual batch.

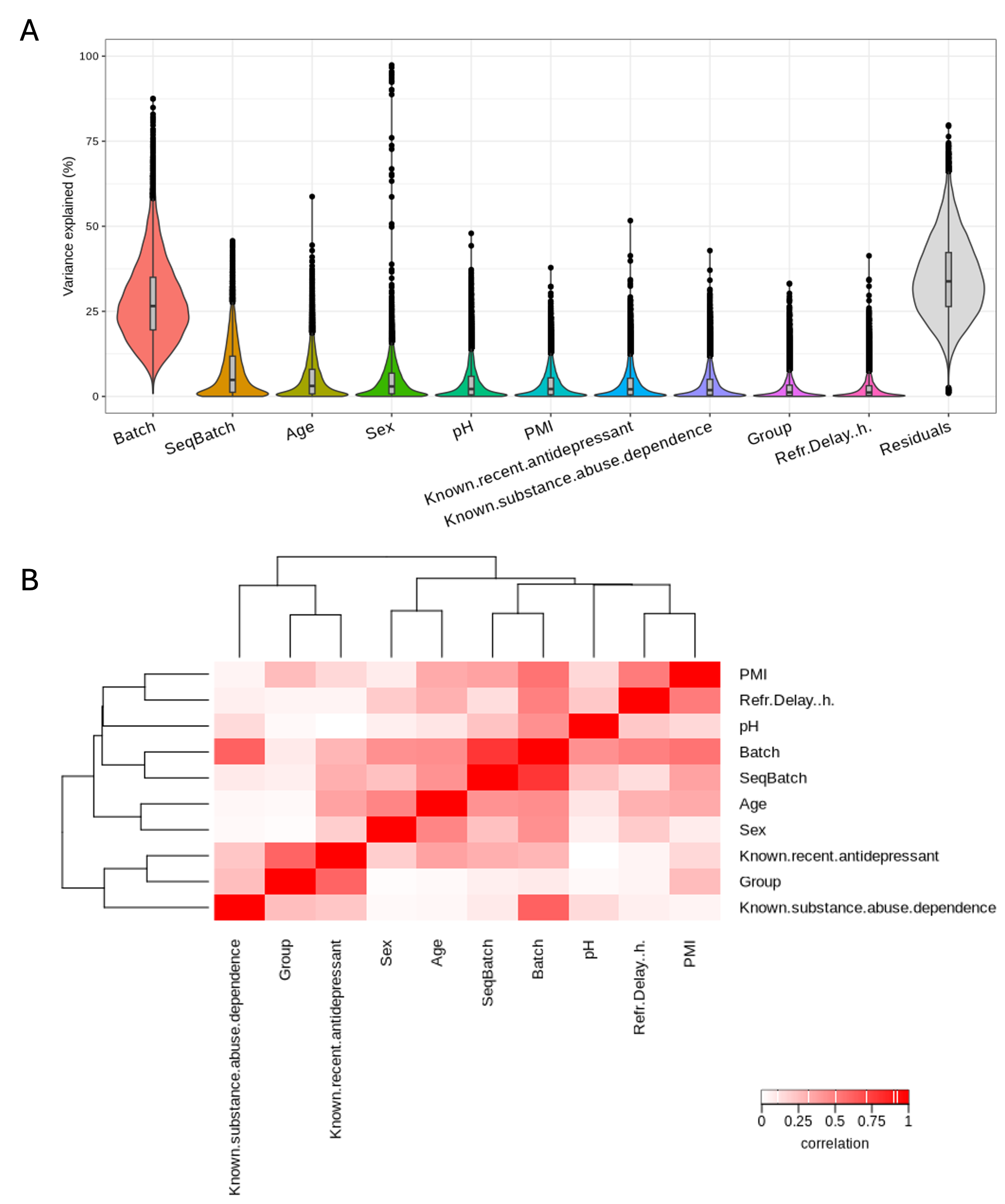

Supplementary Figure 4 – Variance partitioning analysis for selection of covariates. A) Violin plots showing the percent variance explained by each covariate, as well as the residuals, as determined the variance partitioning analysis. The variables on the x-axis are ordered by decreasing variance, with the exception of the residuals. B) Heatmap from canonical correlation analysis (CCA) showing correlations between covariates. The color bar represents the correlation coefficient.

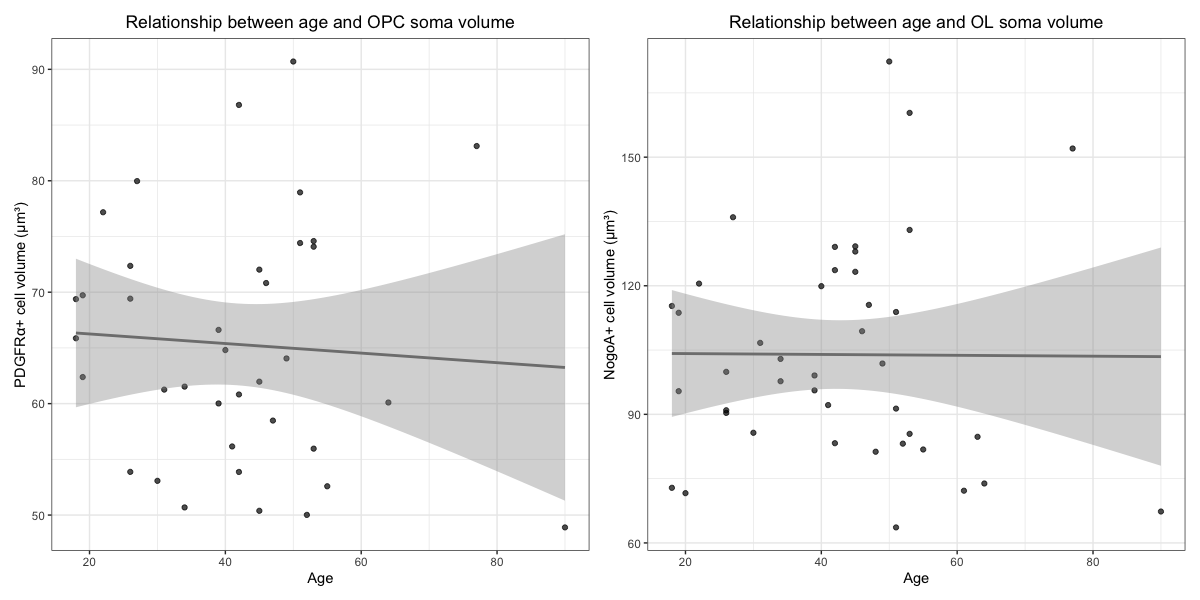

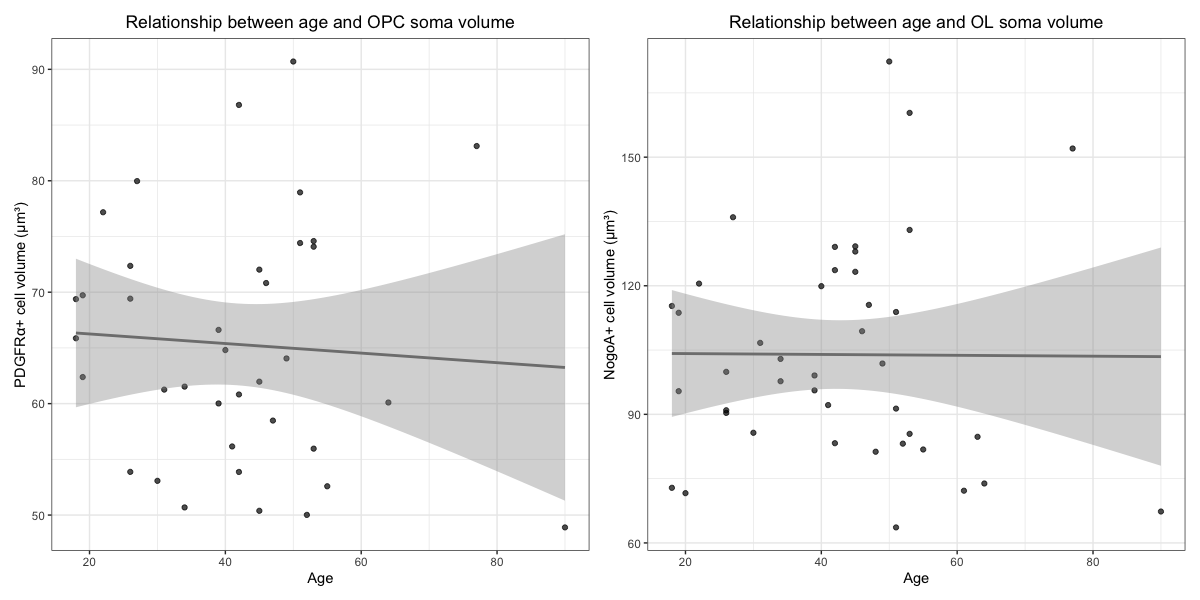

Supplementary Figure 5 - Scatter plots with trend line showing the relationships between age and OL soma volume (left) and age and OPC soma volume (right). Neither OPC nor OL soma volume shows a significant relationship with age.

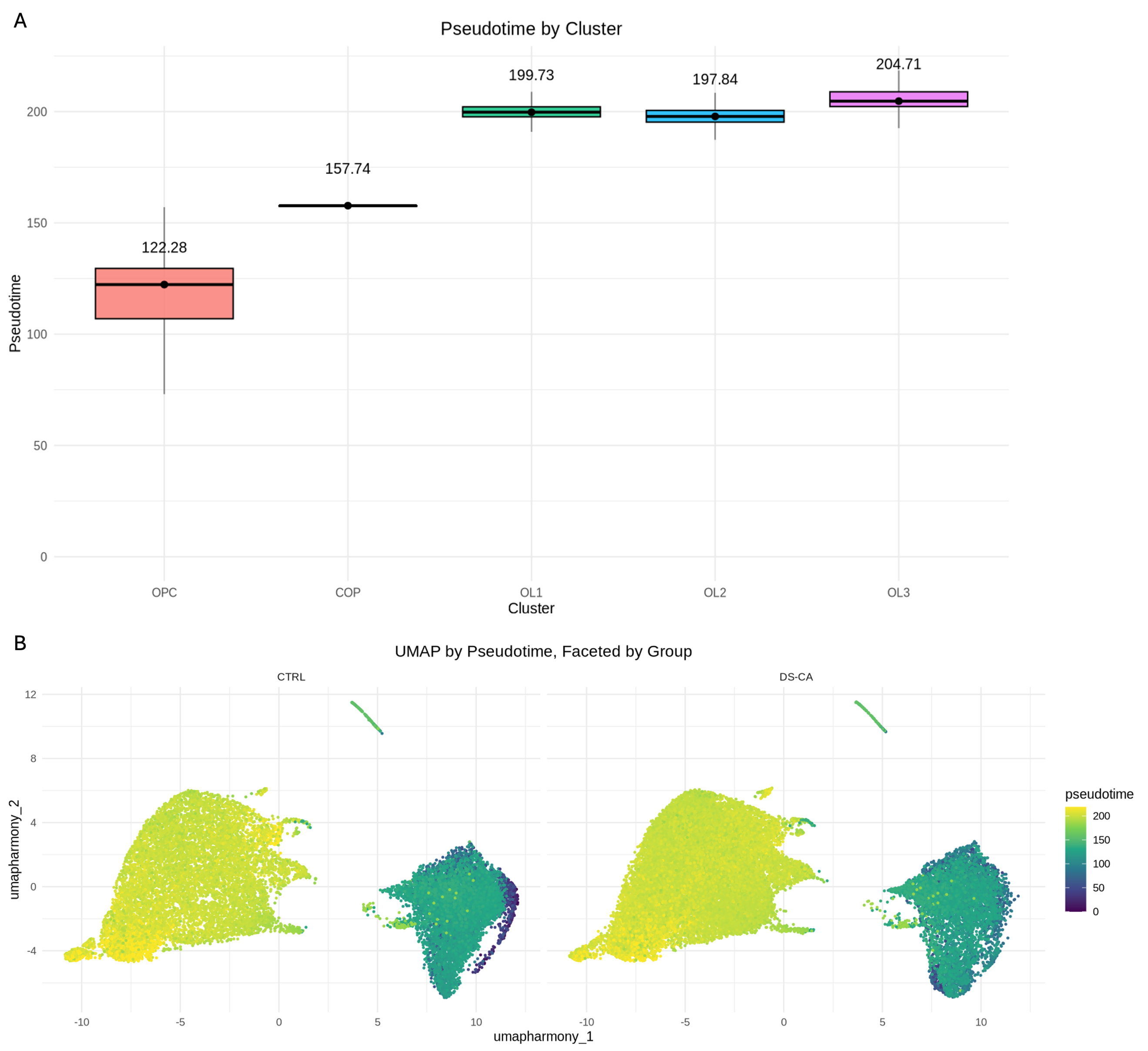

Supplementary Figure 6– A) Box and whisker plot of median pseudotime per OL-lineage cluster. The black dot in the middle of the box represents the median, the value of which is listed on top of the box. The bottom and top of the box represent the first (Q1) and third (Q3) quartiles, respectively. The lower and upper whiskers stretch to the data points that are at least Q1 − 1.5 times the interquartile range and at most Q3 + 1.5 times the interquartile range. B) UMAP of OL-lineage cells OL, colored by pseudotime, and faceted by group (left: CTRL, right: DS-CA). OPC are on the right of each plot, with COP in the middle and OL1, OL2, and OL3 on the left of each plot.

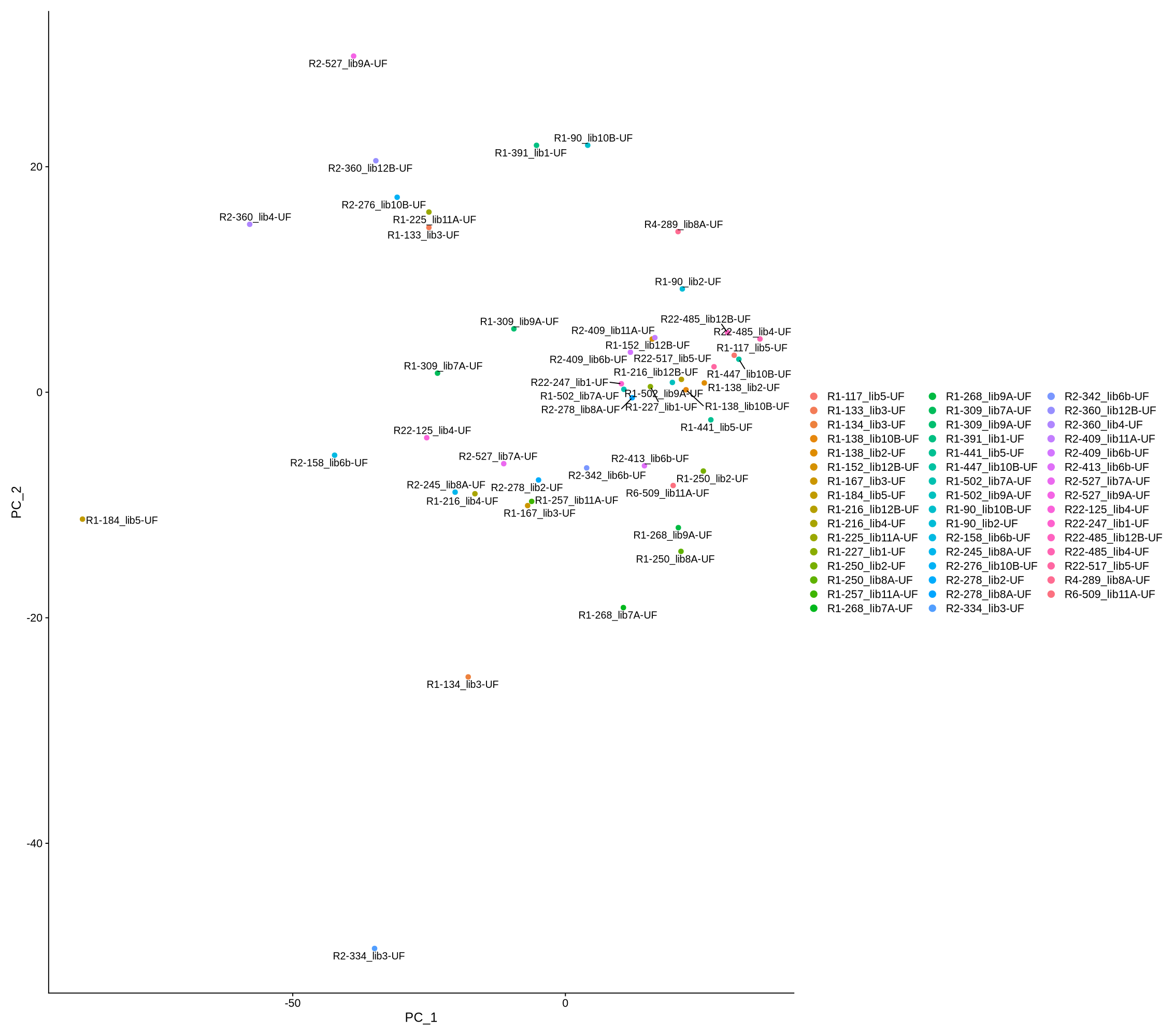

Supplementary Figure 7 – PCA plot for quality control of pseudobulked samples. The points are colored by name of the subject along with the batch it derived from (subject x batch). Most subjects are only in a single batch and as such only one have value. There are no drastic outliers and the subjects that were in two batches are relatively close to each other along these PC dimensions (e.g., R2_360).

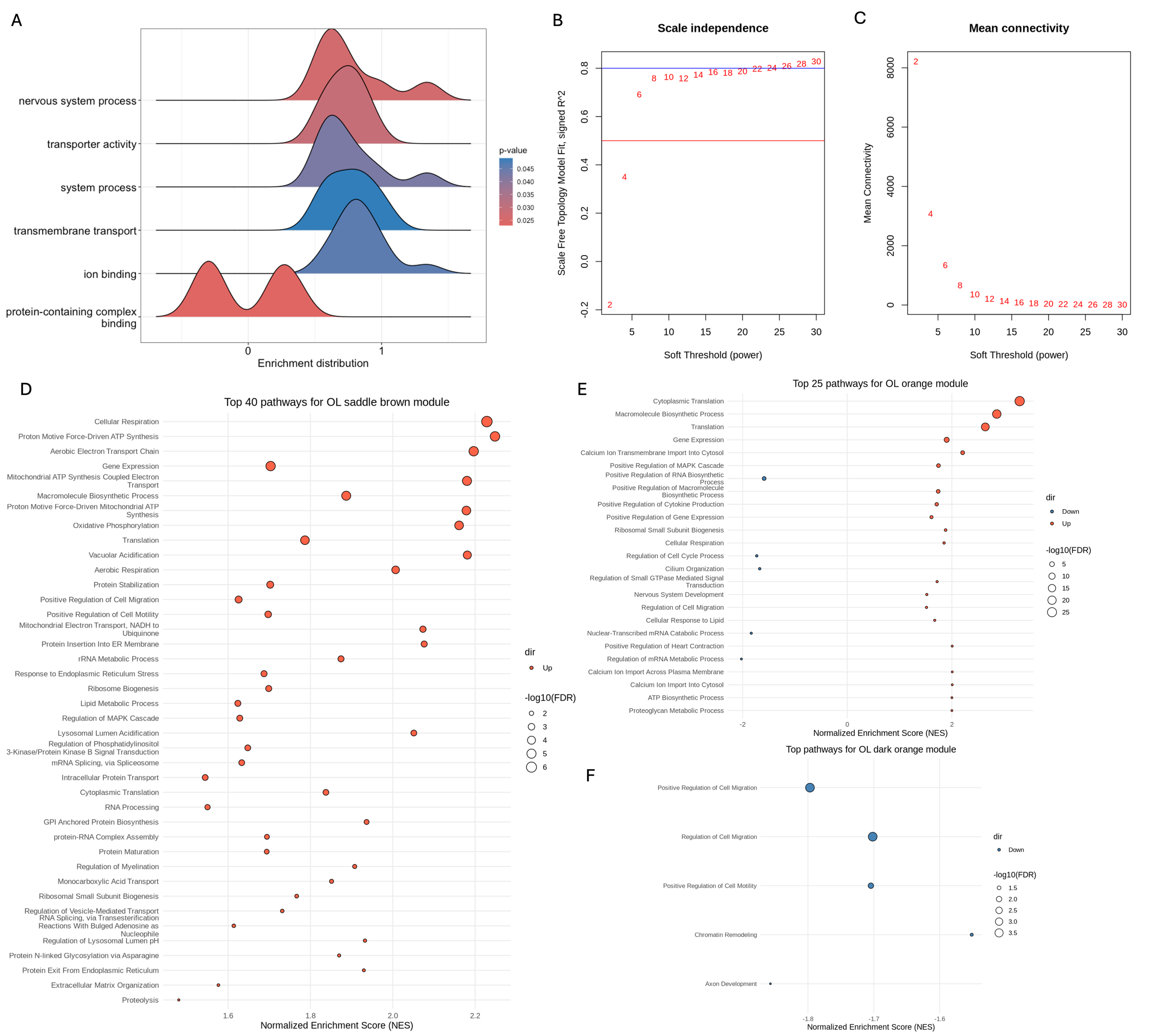

Supplementary Figure 8 –A) Ridge plot showing gene set enrichment of age DEGs in the broad clusters considered together. The x-axis represents the enrichment distribution with negative values indicating a decrease with age and positive values representing an increase with age. The y-axis lists the gene sets enriched in the aging DEGs. The color of the ridge plot represents the p-value. B) Diagnostic plot to select optimal soft thresholding power for WGCNA analysis of OL broad cluster. A soft power of 16 was selected. Plots demonstrate scale independence (left) and mean connectivity (right). On the x-axis of both plots are the soft powers being tested. The y-axis of the scale independence plot refers to the goodness of fit of the gene co-expression network’s degree distribution (distribution of number of connections per gene) to a scale-free distribution (in which the degree distribution follows a power law). A scale free network tends to have a small number of very highly connected (high degree) nodes, and many nodes with few connections (low degree). A R^2^ cutoff of 0.8 (blue horizontal line) is generally considered a useful threshold. The mean connectivity plot y-axis refers to the average sum of edge weights per node (gene). With increasing power, the network becomes sparser as high connectivity nodes (“hubs”) are emphasized. A soft power of 16 was selected due to its R^2^ value approaching 0.8 and its plateaued mean connectivity value. C) Dot plot showing top 40 enriched biological pathways for the genes in the OL saddle brown module, the expression of which is decreasing with age (p = 0.001). Normalized enrichment scores (NES) are represented on the x-axis and gene set names on the y-axis. The size of the circle represents the -log10(false discovery rate) value, with larger dots having lower FDR values. The color of the dot represents the direction of regulation of each term (salmon = up). D) Dot plot showing top 25 enriched biological pathways for the genes in the OL orange module, the expression of which is significantly decreasing with both age (p = 0.04) and group (p = 0.02). Normalized enrichment scores (NES) are represented on the x-axis and gene set names on the y-axis. The size of the circle represents the -log10(false discovery rate) value, with larger dots having lower FDR values. The color of the dot represents the direction of regulation of each term (blue = down, salmon = up). Some terms (e.g. positive regulation of heart contraction) are not relevant here. E) Dot plot showing the enriched biological pathways for the genes in the OL dark orange module, the expression of which is significantly increasing with group (p=0.04) and approaching significance with age (p=0.06). Only these 5 terms were enriched in the parameters set for gsea. Normalized enrichment scores (NES) are represented on the x-axis and gene set names on the y-axis. The size of the circle represents the -log10(false discovery rate) value, with larger dots having lower FDR values. The color of the dot represents the direction of regulation of each term (blue = down).

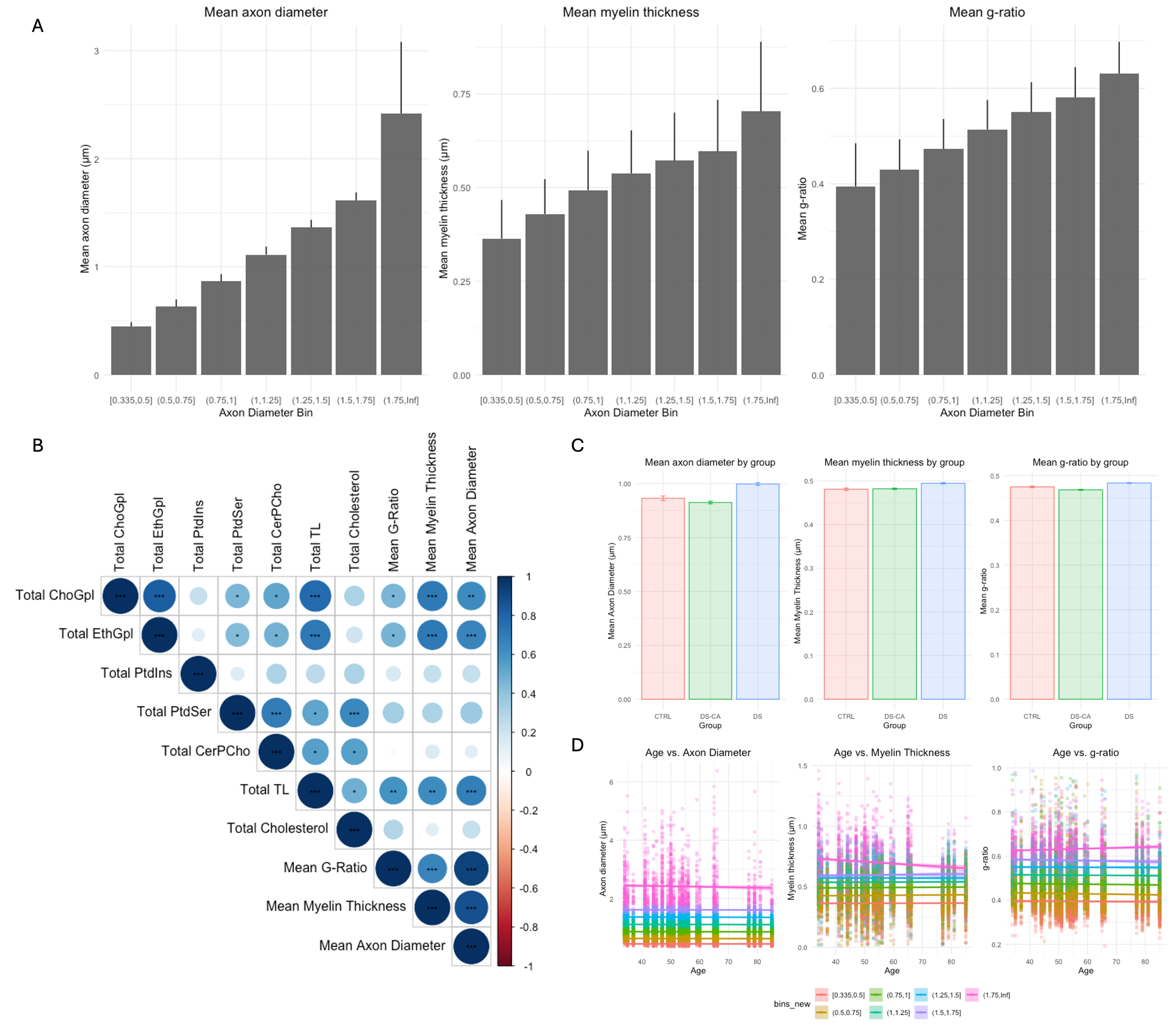

Supplementary Figure 9 – A) Bar plots showing mean + standard deviation of axon diameter (left), myelin thickness (middle), and g-ratio (right) across all 18,822 fibers, segregated by axon diameter bin. B) Correlation plot showing Pearson’s correlations between total lipid concentration and ultrastructure metrics. The significance stars *, **, and *** correspond to the adjusted p ≤ 0.05, 0.01, 0.001, respectively. The color and size of the circles represent the correlation coefficient. C) Bar plots showing mean ± standard error of the mean for axon diameter (left), myelin thickness (middle), and g-ratio (right) by group. D) Scatter plots with trend lines showing the relationship between age and axon diameter (left), myelin thickness (middle), and g-ratio (right). The color of the dots and lines represent the axon diameter bin. ChoGpl: choline glycerophospholipid, EtnGpl: ethanolamine glycerophospholipid, PtdIns: phosphatidylinositol, PtdSer: phosphatidylserine, CerPCho: sphingomyelin, TL: total lipid.

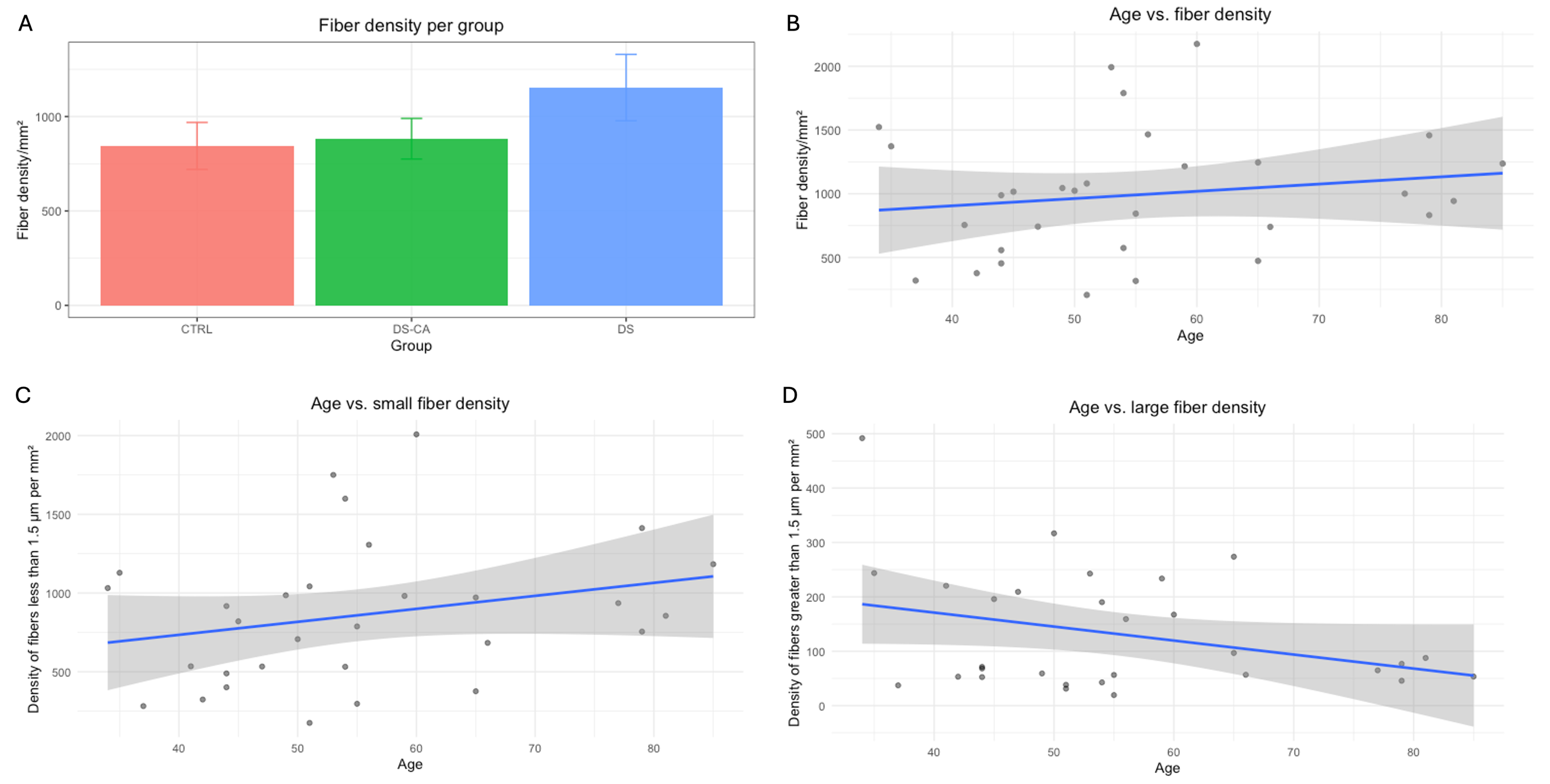

Supplementary Figure 10 – Fiber density is not significantly associated with group or age. A): Bar plot showing mean fiber density plus standard error of the mean per group. No statistically significant difference is observed (p = 0.46). B): Scatter plot with trend line (blue) demonstrating no statistically significant relationship between fiber density and age (p = 0.42). Scatter plot with trend line (blue) showing the relationship between age and fiber density for small (C) and large (D) fiber diameter densities. Small fibers are defined as axons with diameter < 1.5 µm, and large fibers are defined as axons >= 1.5 µm. The density of large fibers appears to decrease with age, while the density of small fibers appears to increase with age.
